# Time-resolved functional connectivity during visuomotor graph learning

**DOI:** 10.1101/2024.07.04.602005

**Authors:** Sophie Loman, Lorenzo Caciagli, Shubhankar P. Patankar, Ari E. Kahn, Karol P. Szymula, Nathaniel Nyema, Dani S. Bassett

## Abstract

Humans naturally attend to patterns that emerge in our perceptual environments, building mental models that allow future experiences to be processed more effectively and efficiently. Perceptual events and statistical relations can be represented as nodes and edges in a graph. Recent work in graph learning has shown that human behavior is sensitive to graph topology, but little is known about how that topology might elicit distinct neural responses during learning. Here, we address this knowledge gap by applying time-resolved network analyses to fMRI data collected during a visuomotor graph learning task. We assess neural signatures of learning on two types of structures: modular and non-modular lattice graphs. We find that task performance is supported by a highly flexible visual system, relatively stable brain-wide community structure, cohesiveness within the dorsal attention, limbic, default mode, and subcortical systems, and an increasing degree of integration between the visual and ventral attention systems. Additionally, we find that the time-resolved connectivity of the limbic, default mode, temporoparietal, and subcortical systems is associated with enhanced performance on modular graphs but not on lattice-like graphs. These findings provide evidence for the differential processing of statistical patterns with distinct underlying graph topologies. Our work highlights the similarities between the neural correlates of graph learning and those of statistical learning.

## 2 Introduction

Spatial and temporal statistical regularities are ubiquitous in our environments. They structure our experiences [1] and inform our expectations about upcoming events [2]. The process of accumulating and applying knowledge of these regularities is known as statistical learning. Operating over a variety of sensory modalities [3], statistical learning allows for the detection of regularities in visual [4, 5, 6], tactile [6], non-linguistic auditory [7, 6], and linguistic [8, 9] sequences. Statistical learning is commonly studied by exposing participants to a sequence of stimuli wherein the probability with which one event follows another, known as the *transition probability*, varies across stimulus pairs [10]. Learning in participants is measured through recognition or recall assessments or by evaluating the relationship between reaction times and transition probabilities [11, 12, 13]. Yet, even in studies demonstrating behavioral evidence for learning, participants generally cannot describe the precise statistical structure underlying the different stimuli, suggesting that the learning process is implicit [14, 15, 16, 17].

The breadth of evidence that supports statistical learning from across multiple modalities, age groups [18, 19, 20, 21], and species [22, 23], and its putative role in cognitive function [8] has motivated investigations into its underlying neural bases. Research often considers the persistence of these neural bases across modalities and their relationships to other forms of associative and implicit learning [3]. Electrophysiological and fMRI studies have implicated several regions in various aspects of statistical learning, including the superior temporal gyrus [10, 24, 25], the inferior frontal gyrus [10, 26, 27], the inferior temporal cortex [28, 29, 30, 13, 10], and the medial temporal lobe [13, 31, 32, 33, 34]. Although the proposed relationships between activity in these areas and statistical learning vary from study to study, most relationships fall into one of three broad categories: (1) increased activity is associated with training or testing on structured sequences compared to unstructured sequences [10, 24, 25, 27, 13]; (2) increased activity is associated with enhanced performance on structured sequences [10, 26]; (3) patterns of activity are similar for statistically related stimuli [28, 29, 30, 31].

Previous research on statistical learning has focused primarily on how humans learn transition probabilities between temporally adjacent stimuli. This work has paved the way for research examining how the brain processes more complex statistical structures [35]. Questions in this domain pertain to so-called *structure learning* [36, 37], *or more specifically, graph learning* [38]. In this form of learning, sequences are generated via walks on graphs composed of stimulus events (nodes) and event-to-event transitions (edges). Graphs capture the complexity of a wide range of real-world information, such as speech, music, and social behavior, where large numbers of possible events exhibit inter-dependencies in organized, hierarchical relationships [39]. Recent work has also shown that the organization (or *topology*) of transition graphs can influence behavior [40, 41, 42, 43, 44] and neural activity [35, 45, 46, 47, 48]. In particular, when a graph is characterized by modules (densely inter-connected groups of nodes), response times for between-module transitions are higher than those for within-module transitions [35, 41, 42, 44]. Moreover, neural representations for items within the same module are markedly similar in the inferior frontal gyrus, superior temporal gyrus, anterior temporal lobe, and hippocampus [35, 45]. Modular graphs yield consistent higher-dimensional representations compared to non-modular graphs [46]. The topology of the transition graph also influences the accuracy with which the identity of visual stimuli can be decoded from neural representations in the lateral occipital cortex. Participants who learn from modular graphs exhibit higher decoding accuracy than those learning from non-modular lattice graphs. Furthermore, accuracy also correlates with variations in static functional connectivity among the medial temporal lobe, lateral occipital cortex, and the postcentral gyrus during learning [46]. Taken together, these findings suggest that neural representations of stimuli are shaped by the topology of the underlying statistical structure and that the strength of such representations is associated with differences in inter-areal communication.

Despite significant recent progress in graph learning research, several questions remain unexplored. First, prior work has predominantly focused on transient patterns of activation associated with individual edges or nodes in a single graph type. It remains unclear, however, whether activity observed throughout learning differs based on topology. Second, prior work has centered itself on univariate measures of regional activity or static measures of inter-regional connectivity, with limited emphasis on dynamic interactions [3, 49]. The former are often obtained by averaging functional connections throughout a learning task, and, therefore, finegrained functional information is lost. Given these gaps, little is known about the dynamic reorganization of inter-areal functional connections that likely underlies graph learning, as has been observed for other learning and working memory task domains [50, 51, 52, 53]. Here, we address these gaps directly by performing a time-resolved functional connectivity analysis of fMRI data collected in 31 healthy individuals during a visuomotor graph learning task. We compare the learnability of two types of graphs: modular and non-modular lattice graphs [43, 46]. We test the following two hypotheses: (1) brain networks that support graph learning include regions that overlap with those implicated in statistical learning, and (2) task performance is supported by different brain networks depending on the presence or absence of modularity. By testing these hypotheses, we contribute to an improved understanding of the neural processes associated with statistical learning.

## 3 Results

### 3.1 Task

Thirty-four right-handed individuals (aged 18–34 years, 68% female) underwent nine fMRI scans: four during rest and five while performing a visuomotor graph learning task designed to compare the learnability of modular and lattice graphs [46, 43]. Due to exclusions for poor task performance and technical challenges in recording behavioral data, our analyses are performed on a total of 31 participants. Visual stimuli consisted of 15 unique abstract shapes (Fig. 1a, *left*). The corresponding motor responses comprised the set of all 15 possible one-or two-button finger presses on a five-button controller (Fig. 1a, *right*). Participants were randomly assigned to one of two graphs, modular or lattice, where each graph consisted of 15 nodes and 30 undirected and unweighted edges (Fig. 1b, *left*). We generated sequences of visual stimuli by taking a random walk on each participant’s assigned graph structure; each node in the sequence was a single shape-response pair, and each edge was a valid transition between two stimuli (Fig. 1b, *right*). The three-way shape-response-node associations were randomized and did not follow any organizational rules. Both the modular and lattice graphs were regular, meaning that each node had the same number of neighbors. Thus, the transition probabilities were uniform across all pairs of connected nodes. However, in the modular graph, nodes in the same module were more likely to follow one another in a random walk than nodes in different modules since distinct modules were connected only by a single edge. The broader goal was for participants to learn the statistical dependencies between shape-response pairs as dictated by the placement of edges in the underlying graphs. Therefore, shape-response-node associations were mapped one-to-one and were consistent within individuals but varied across them.

**Figure 1.**
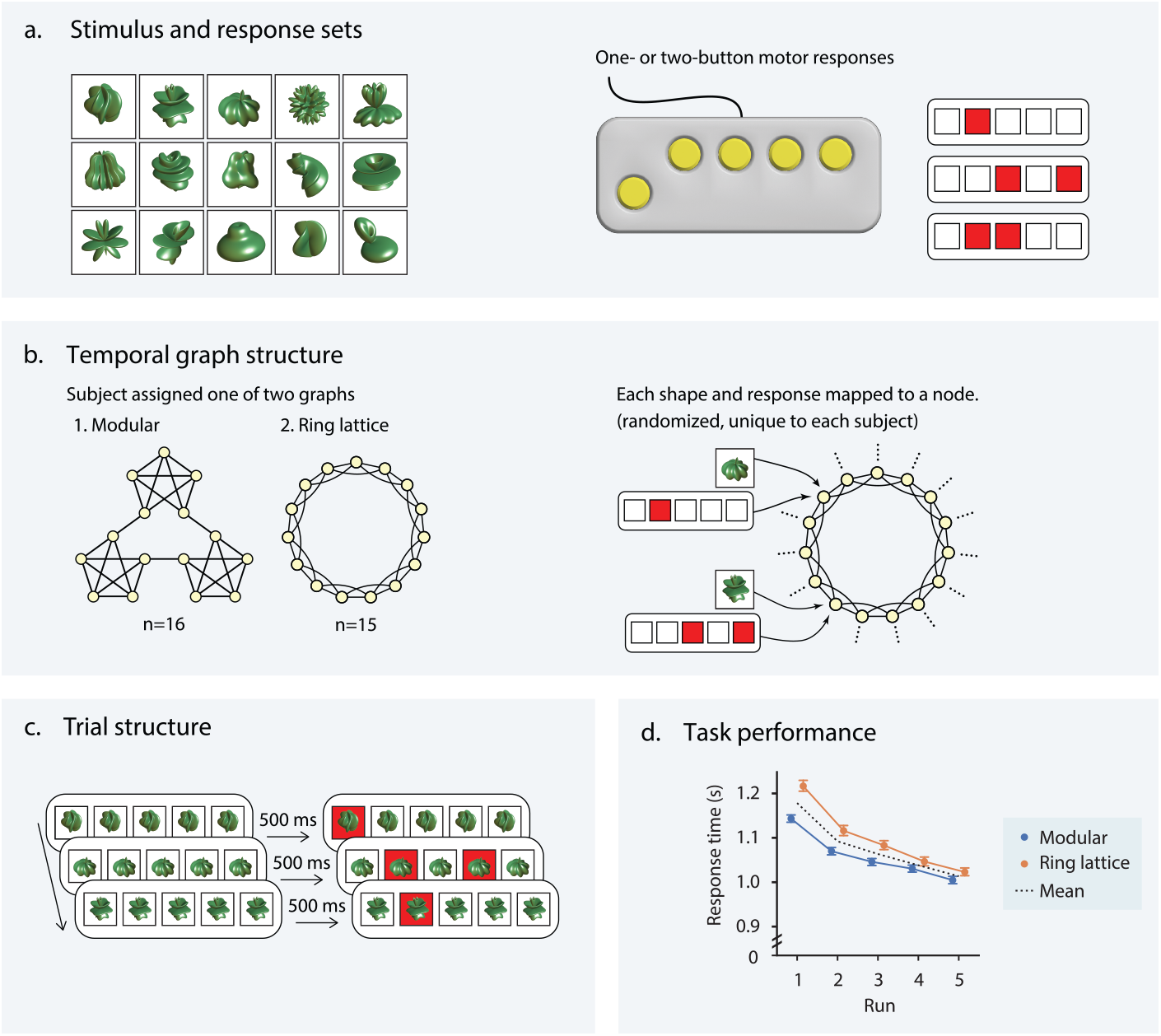
Visuomotor graph learning task. (a) During the visuomotor learning task, study participants were trained on a set of 15 abstract shapes (*left*) and 15 possible one- or two-button combinations on a controller (*right*). (b) Each participant was assigned to one of two conditions: stimuli organized on a modular graph and stimuli organized on a ring lattice graph (*left*). The 15 shapes and motor responses were mapped to one of the 15 nodes in the assigned graph (*right*). To control for differences among shapes and motor responses, each mapping was random and unique to a participant. (c) Participants were visually instructed as to which buttons to press (i.e., those indicated by the red squares). To encourage participants to learn the shape-motor response mappings, the shape appeared 500 milliseconds before the motor command. (d) Response time (seconds) as a function of task run for correct trials; color indicates whether participants were trained on the lattice graph (orange) or the modular graph (blue); colored markers indicate participant averages for each run; error bars indicate 95% confidence intervals; black line indicates mean across both graph groups. Adapted with permission from [46].

To demonstrate that participants successfully learned statistical dependencies between stimuli as encoded in their assigned graphs, Kahn et al. [46] examined their response times over the five task runs. The reaction time between a stimulus and the associated response is influenced by one’s expectations of the current stimulus, given the preceding stimulus; if a participant has learned the statistical likelihood with which one shape (or, equivalently, one motor response) follows another—in other words, if they have learned the structure of the underlying graph—they can improve their predictions about incoming stimuli and, in turn, respond more quickly. Averaging response times across all correct trials in each run, Kahn et al. [46] showed that response times decreased as a function of the run for participants in both the lattice and modular groups (Fig. 1d), indicating continuous learning of underlying graph structures.

### 3.2 Network analysis

We seek to investigate the relationship between correlations in neural activity and graph learning. Therefore, we begin by conducting a whole-brain functional connectivity analysis of fMRI data collected during the task and rest conditions (Fig. 2a). Using the Schaefer 300-parcel 17-system cortical atlas [54], the corresponding Buckner 17-parcel cerebellar atlas [55], and the Scale II, 3T version of the 32-parcel Melbourne subcortical atlas [56], we parcellate the brain into 349 regions and extract the average BOLD time series for each region (Fig. 2b). Defining the strength of a functional connection as the product-moment correlation between time series, we first compute static functional connectivity matrices for each scan by using the full length of each time series to calculate connection strengths.

**Figure 2.**
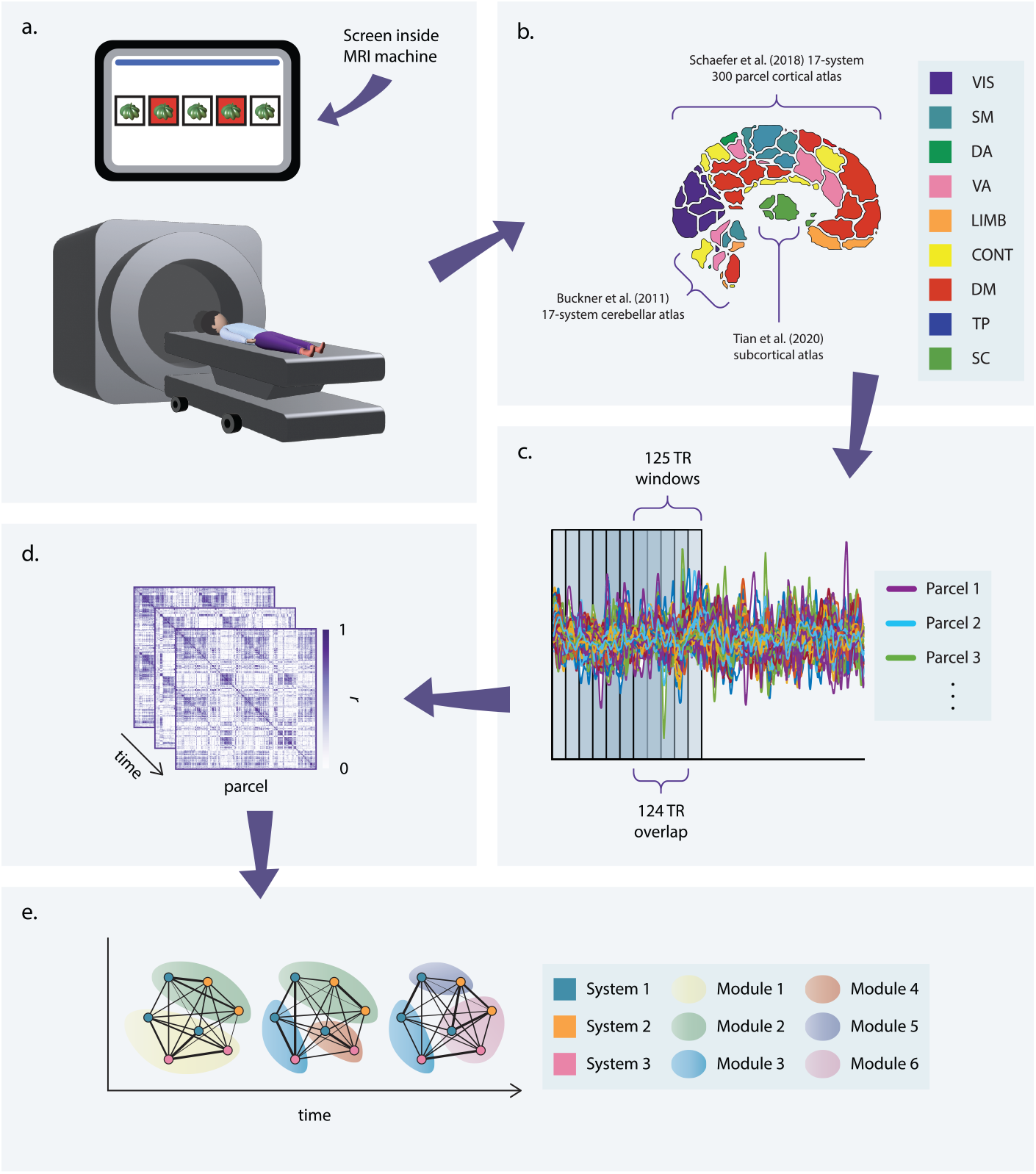
Study design. (a) Participants completed the visuomotor graph learning task while in an MRI scanner. (b) To parse whole-brain fMRI data into regions of interest, we applied 3 parcellation schemes: a 17-system, 300-parcel cortical atlas [54]; a 17-parcel cerebellar atlas, provided with an allocation scheme of each parcel to one of the 17 cortical systems [55]; and a 32-parcel subcortical atlas [56]. Averaging over subsystems of the 17-system atlases (e.g., over visual systems A and B) and across all subcortical parcels yielded a coarse-grained 9-system parcellation. (c) Time series were extracted for each parcel. To obtain time-resolved measures of network structure, these time series were divided into overlapping windows. (d) Functional connectivity matrices were obtained for each window by computing the product-moment correlation coefficient, *r*), for each pair of time series in a given window. (e) The functional connectivity matrices from panel (d) encode edge weights for a series of fully connected networks for each scan. Community detection was performed on each network to identify modules corresponding to groups of strongly connected nodes.

Pursuant to the characterization of more fine-grained changes in functional connectivity during the learning process, we then parsed each time series into sliding windows (Fig. 2c) and computed functional connectivity matrices for each window (Fig. 2d), thereby generating time-resolved functional networks. We generated four such networks for each scan by varying the degree of overlap between windows; statistics reported here are associated with networks generated using maximal overlap (see Section 5.2.1), though only results that were statistically significant for all four networks are included. For the time-resolved analyses, we elected to investigate community structure, as time-resolved measures of community structure have previously been associated with learning and memory [51, 57, 50, 52, 58, 53]. To this end, we applied a generalized Louvain-like community detection algorithm to the time-resolved networks to obtain community assignments for each parcel in each window (Fig. 2e). We then summarized these results in module allegiance matrices, wherein each element is the frequency with which two brain regions are assigned to the same community over time windows for a given network.

We calculated three additional metrics from the module allegiance matrices: flexibility, recruitment, and integration. To determine the stability of community assignments, we calculated flexibility or the frequency with which a region switches communities over time windows. To estimate the degree to which regions within the same cognitive system interact, we calculated recruitment, or the probability with which a given region is assigned to the same community as the other regions in its cognitive system over time windows. Finally, to assess inter-system interactions, we calculated integration, or the probability with which a pair of regions from different cognitive systems is assigned to the same community over time. We considered the following eight cortical systems, as defined by the Schaefer atlas: visual, somatomotor, dorsal attention, ventral attention, limbic, frontoparietal control, default mode, and temporoparietal. We also included one subcortical system encompassing the hippocampus, thalamus, amygdala, caudate nucleus, nucleus accumbens, putamen, and globus pallidus.

Static functional connectivity, flexibility, recruitment, and integration—henceforth referred to collectively as “network metrics”— were first examined on a global level, averaging across parcels, then at a system level, averaging across parcels within each of the nine systems, and finally at a parcel level. For the two metrics associated with pairs of regions, static functional connectivity and integration, intra-system values were excluded from statistical analysis; for example, if two parcels belonged to the visual system, then their static functional connectivity value was excluded from the global- and system level averages, and any associated *p*-values were excluded from correction for multiple comparisons in parcel level analyses.

We began by comparing rest and task data using paired *t*-tests, averaging across runs and parcels to obtain one static functional connectivity, flexibility, recruitment, and integration value for each participant. We then repeated this comparison for each system (in the case of flexibility and recruitment) or pair of systems (in the case of static functional connectivity and integration). We do not include parcel-level analyses for rest-task comparisons due to the volume of significant results. The *p*-values associated with the paired *t*-tests were Bonferroni-corrected for multiple comparisons within each scale and network metric. Next, to assess the effects of task run (one to five) and graph type (lattice versus modular) on functional network architecture, we fit network metrics computed from task networks to multilevel mixed-effects models with fixed effects of graph, task run, and the interaction between the two, and one random effect corresponding to the participant. We generated a new model for each metric and each system or pair of systems and then each parcel or pair of parcels. Finally, to determine whether individual variability in task performance could be accounted for by differences in functional network architecture, and to what degree this relationship was modulated by task run and graph type, we fit reaction time data to models with fixed effects of graph, task run, and network metric, and the two- and three-way interactions between these parameters, and one random effect for the participant, again generating a new model for each metric and each system or pair of systems and then each parcel or pair of parcels. The *p*-values for fixed effects were calculated via the Satterthwaite approximation and, within each model type, scale, and network metric, corrected for multiple comparisons using the false-discovery rate procedure.

### 3.3 Functional brain network organization during task and rest

To capture task-related connectivity patterns, we examined the difference in static functional connectivity and time-resolved community structure between task and rest conditions. At the global level, we found that static functional connections were, on average, stronger during the task than during rest (*t* = 7.108, *df* = 30, *p*_FDR_ < 0.0001, Cohen’s *d* = 1.277). This trend persisted at the system level; all 36 inter-system connection strengths were higher during the task than during rest (2.756 < *t* < 11.278, *df* = 30, *p*_FDR_ < 0.01, 0.495 < Cohen’s *d*< 2.026; Fig. 3, *left*). These findings suggest that task performance is supported by a higher degree of communication between different brain regions than occurs during rest.

**Figure 3.**
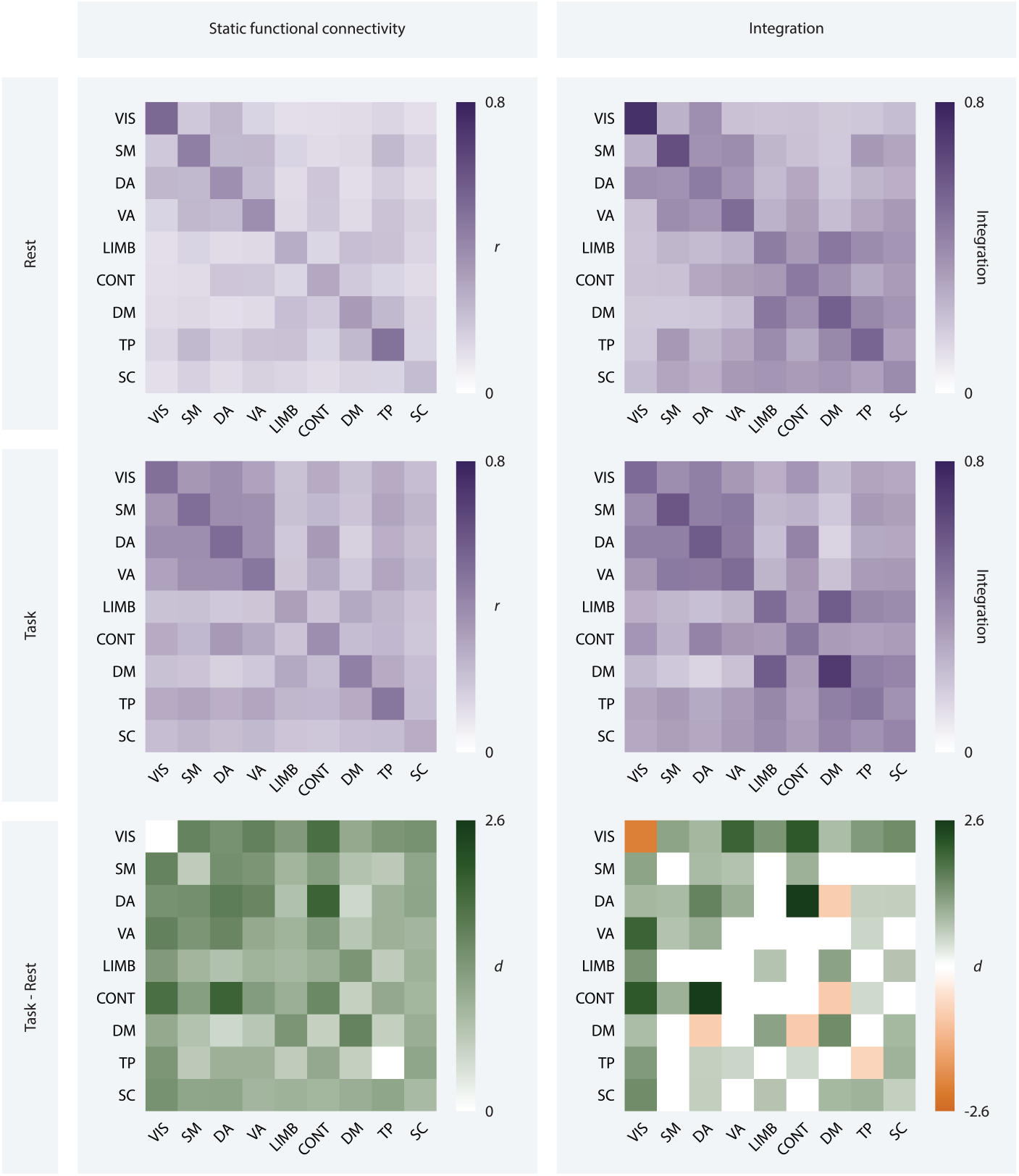
Static and time-resolved metrics of functional organization during rest and task. Static functional connectivity (*left*) and integration (*right*) matrices for resting-state (*top*) and task fMRI data (*middle*), averaged across participants, task runs, and systems. Each entry on the x- and y-axes corresponds to a system. The darker the square, the stronger the connection between the corresponding systems (*left*), or the more frequently they are assigned to the same module (*right*). Effect size matrices (*bottom*) show the magnitude and direction of the effect of condition on static functional connectivity and integration using Cohen’s *d*. The darker the purple or orange, the larger the magnitude of the effect. Purple squares indicate an increase in value from rest to task, while orange squares indicate a decrease in value from rest to task, and white squares indicate no difference surviving a statistical threshold of *p*_FDR_ < 0.05.

Turning to time-resolved community structure, we found that flexibility was lower during task than during rest at the global level (*t* = −3.539, *df* = 30, Cohen’s *d* = −0.636, *p*_FDR_ = 0.001). At the system level (Fig. 4, *top*), the dorsal attention, limbic, frontoparietal control, and default mode systems also displayed reduced flexibility during task performance (−6.175 *< t <* −4.188, *df* = 30, *p*_FDR_ < 0.001, −1.109 < Cohen’s *d <* −0.752), while the visual system showed increased flexibility (*t* = 2.552, *df* = 30, *p*_FDR_ = 0.019, Cohen’s *d* = 0.458). These results indicate that community structure is generally more stable during task performance as compared to rest, but that regions in the visual system interact with many different communities in response to task demands.

**Figure 4.**
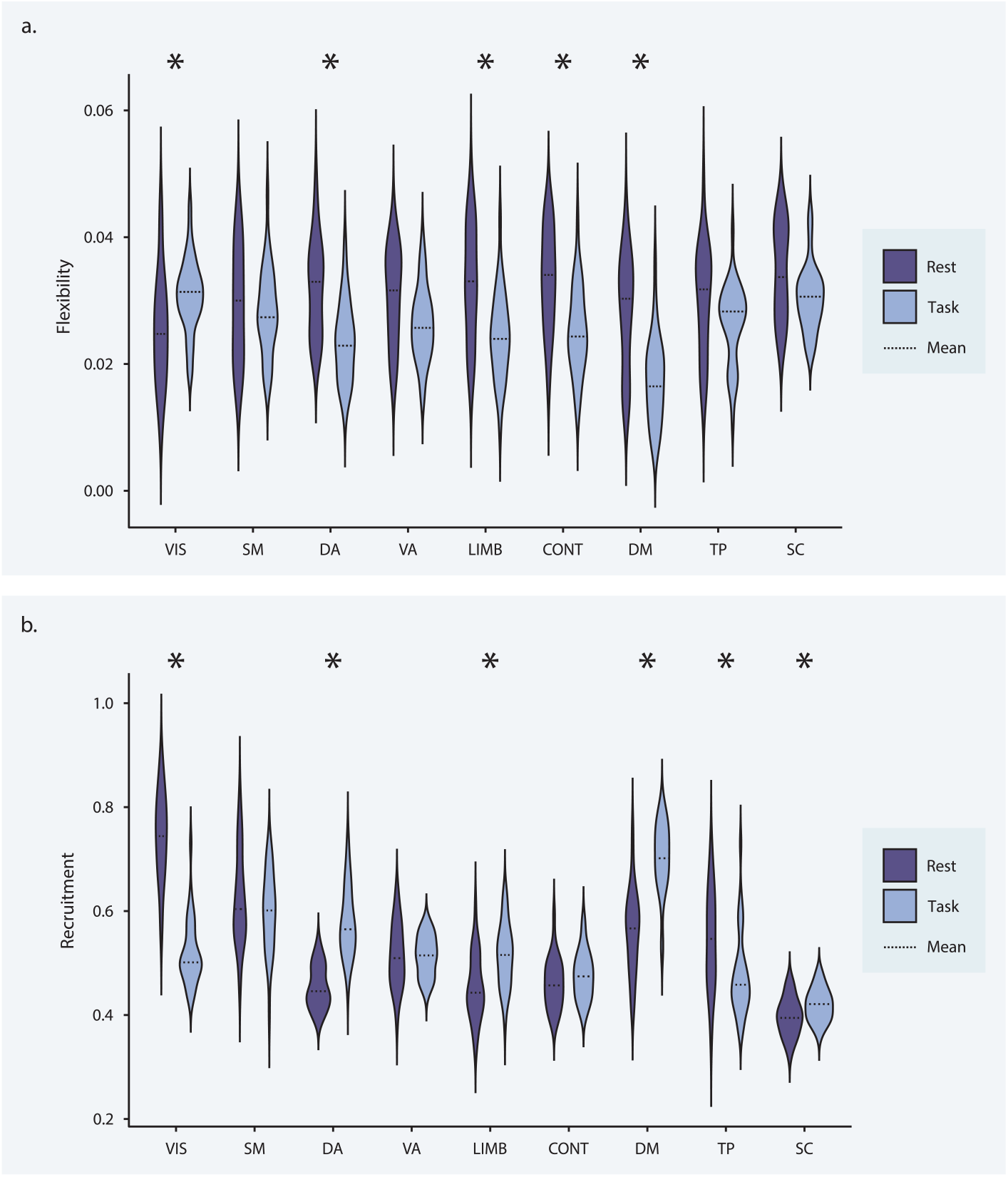
Recruitment and flexibility during rest and task. (a) Average recruitment by system for rest (purple) and task (blue) data. The dashed line indicates the mean. Asterisks indicate differences surviving a statistical threshold of *p*_FDR_ < 0.05. (b) Average flexibility by system for rest (purple) and task (blue) data. The dashed line indicates the mean. Asterisks indicate differences that survive a statistical threshold of *p*_FDR_ < 0.05.

No significant trend in recruitment was found at the global level. At the system level (Fig. 4, *bottom*), dorsal attention, limbic, default mode, and subcortical regions exhibited a higher degree of recruitment during task performance (3.306 < *t* < 8.769, *df* = 30, 0 < *p*_FDR_ < 0.004, 0.594 < Cohen’s *d* < 1.575). Visual and temporoparietal regions showed reduced recruitment during task performance (−12.057 < *t* < −3.928, *df* = 30, *p*_FDR_ < 0.001, −2.166 < Cohen’s *d* < −0.706). Thus, changes in intra-system cohesiveness during task were heterogeneous across systems but tended toward increased cohesiveness. There was, however, a notable functional dissolution of the visual system, potentially indicating distinct roles for different visual regions.

On average, integration was higher during task performance than during rest (*t* = 9.334, *df* = 30, Cohen’s *d* = 1.677, *p*_FDR_ < 0.0001). This trend largely persisted at the system level; 21 pairs of systems (58%) were more integrated during task than during rest (2.681 < *t* < 14.379, *df* = 30, *p*_FDR_ < 0.019, 0.481 < Cohen’s *d* < 2.583; Fig. 3), *right*). Effect size was largest for the integration of the dorsal attention system with the frontoparietal control system (*t* = 14.379, *df* = 30, *p*_FDR_ < 0.0001, Cohen’s *d* = 2.583), followed by the integration of the visual system with the frontoparietal control (*t* = 12.115, *df* = 30, *p*_FDR_ < 0.0001, Cohen’s *d* = 2.176) and ventral attention systems (*t* = 11.272, *df* = 30, *p*_FDR_ < 0.0001, Cohen’s *d* = 2.025). Notably, the integration of the visual system with all other systems was greater during task performance.

The integration of the default mode system with the frontoparietal control and dorsal attention systems were the only integration values that were lower during task performance than during rest (−4.98 < *t* < −4.698, *df* = 30, *p*_FDR_ < 0.0001, −0.894 < Cohen’s *d* < −0.844). When compared to community structure during rest, integration during task was markedly higher, indicating an increase in dynamic inter-system interactions. The most prominent changes occurred among visual, attentional, and executive function regions. Visual regions exhibited heightened communication with all other systems, while top-down processing regions interacted more with each other and less with the default mode system.

### 3.4 Functional brain network organization changes over task run

We next sought to determine whether functional brain network organization (1) changed over the course of the five task runs and (2) differed by graph type. We did not find any significant effects of graph type, task run, or the interaction between the two on static functional connectivity, flexibility, recruitment, or integration at the global or system levels. However, at the parcel level, 157 pairs of regions (0.3%) showed significant changes in integration over the five task runs (Fig. 5); 129 of these pairs exhibited increased integration over time (3.641 < *t* < 5.96, 122 < *df* < 151, 0.001 < *p*_FDR_ < 0.05) and 28 exhibited reduced integration over time (−5.036 < *t* < −3.632, *df* = 122, 0.008 < *p*_FDR_ < 0.05). 40% of all pairs for which integration changed over time were comprised of a visual region and a ventral attention region, all of which became more integrated over time. This result likely did not reach significance at the system level as almost all parcels exhibiting this pattern were in the insular, opercular, frontal medial, and parietal medial regions of the ventral attention system (corresponding approximately to ventral attention system A in the 17-system Schaefer atlas) and the right hemisphere visual system. Further, 16% of all pairs for which integration changed over time were comprised of a visual region and a somatomotor region, all of which also became more integrated over time. The majority of pairs that became less integrated over time (75%) were comprised of a frontoparietal control region and either a visual or dorsal attention region. These results indicate that as learning progressed, interactions between regions of the visual and ventral attention systems increased, while frontoparietal control regions became less functionally connected to visual and dorsal attention regions. Broadly, this suggests an increasing influence of bottom-up or stimulus-driven signals on visual processing and a decreasing influence of top-down control signals over task runs.

**Figure 5.**
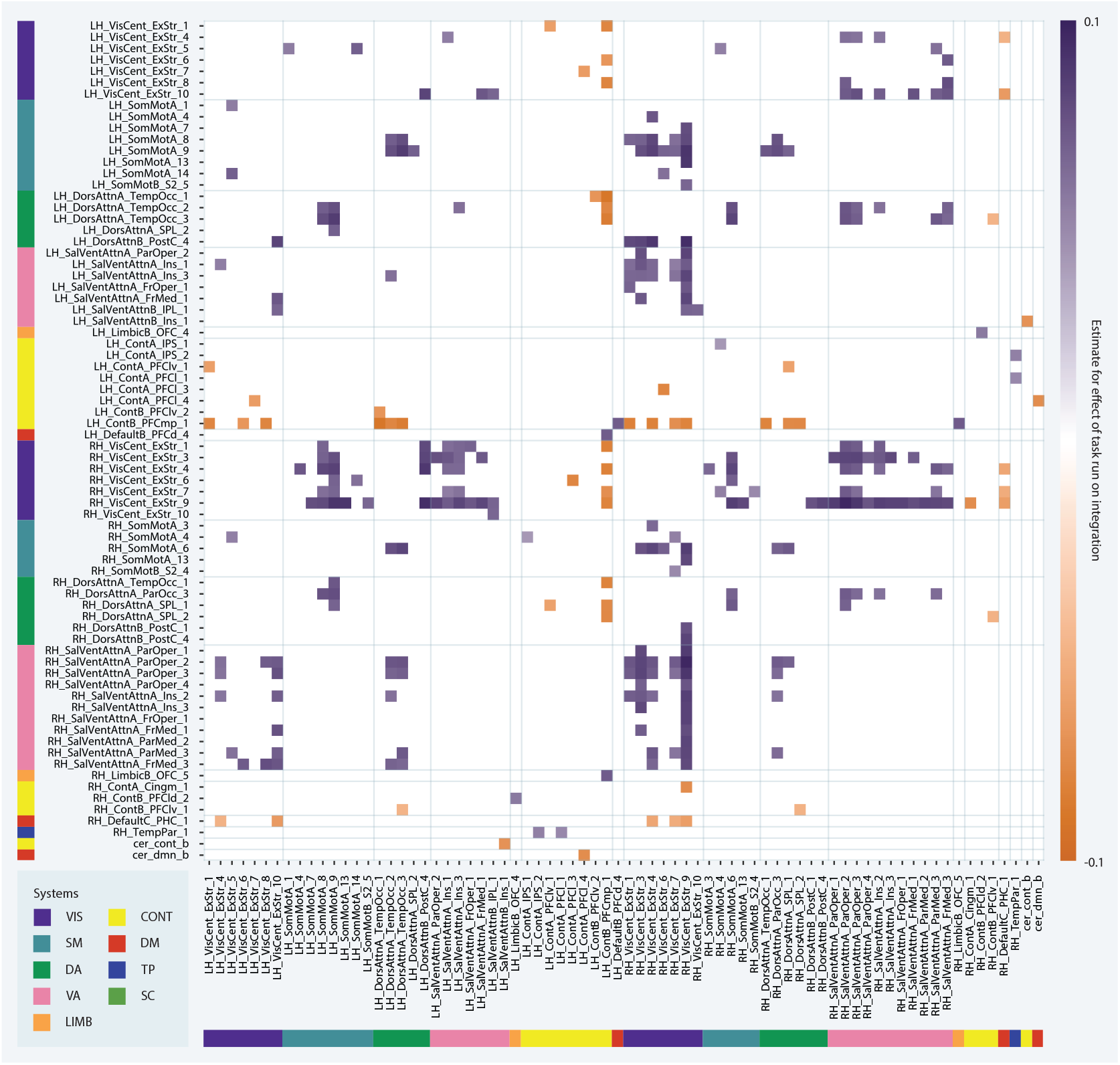
Network changes over task run. Main effect of task run on integration values at the parcel level. Each entry on the x- and y-axes corresponds to a parcel. Each square of the matrix indicates the estimate for the effect of run according to the corresponding multilevel model.

### 3.5 Response time as a function of graph, task run, and functional brain network organization

Finally, we examined the relationship between functional brain network organization and behavior. No models of response time that included static functional connectivity or flexibility as a fixed effect were statistically significant. We found a significant graph *×* recruitment interaction effect at the global level (*t* = −2.388, *df* = 120.4, *p*_FDR_ = 0.018; Fig. 6a), wherein high average levels of recruitment were associated with lower response times for participants trained on the modular graph than for participants trained on the lattice graph. More specifically, we observed a negative relationship between average recruitment and response time for the modular group and a slight positive relationship between average recruitment and response time for the lattice group, indicating that participants who performed better in the modular group tended to exhibit greater system recruitment. A similar trend was found at the system level for the limbic system such that high limbic recruitment was associated with lower response times for participants in the modular group than participants in the lattice group (*t* = −3.289, *df* = 120.02, *p*_FDR_ = 0.012; Fig. 6b). These graph *×* recruitment effects on response time point to an association between the cohesiveness of cognitive systems and successful task performance for the modular group and, intriguingly, the opposite association for the lattice group. In particular, participants in the modular group showed improved performance when limbic regions were working in concert.

**Figure 6.**
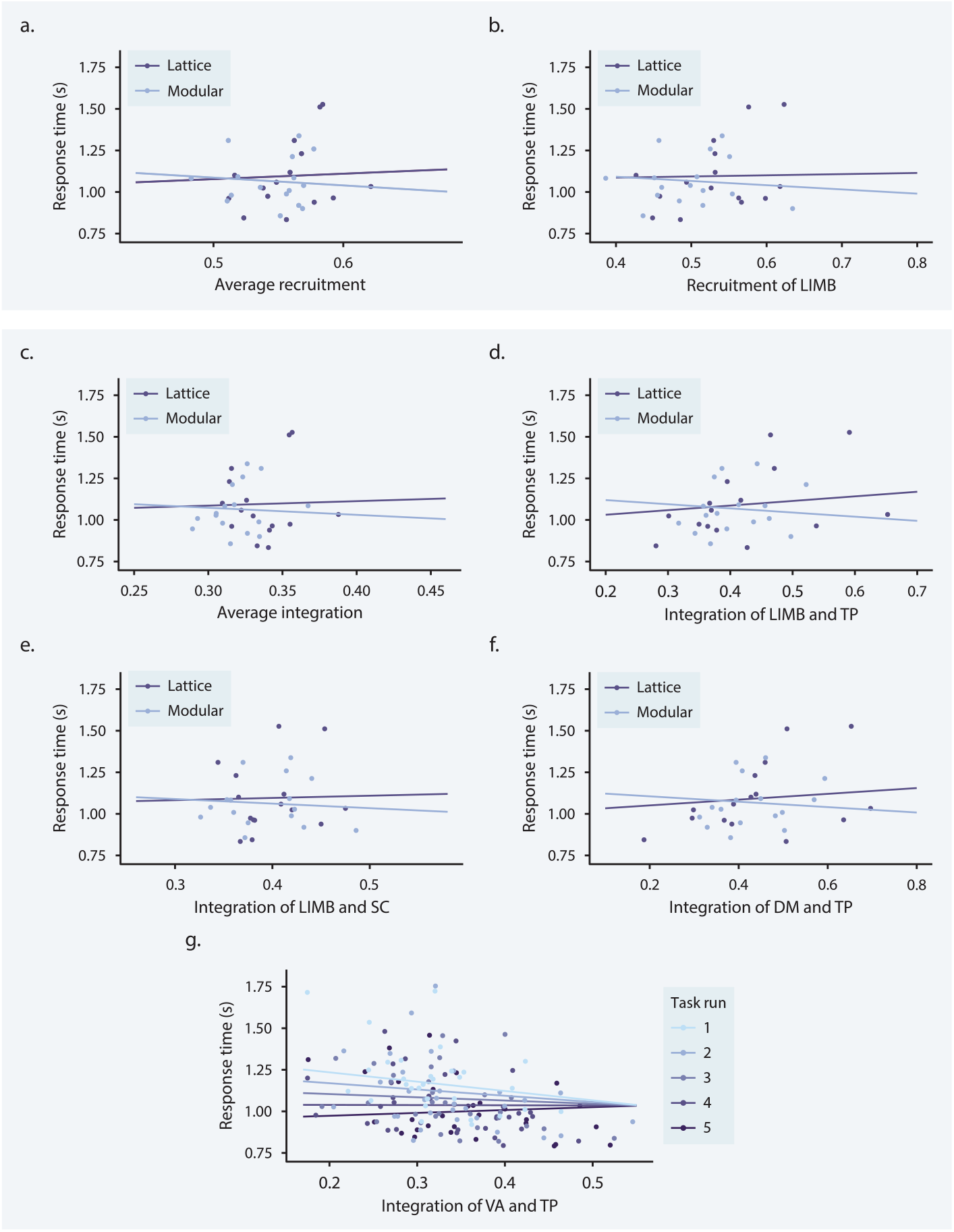
System level network predictors of response time. (a–f) Interaction effect of graph × network metric on response time for three network metrics. Each data point represents a participant; purple and blue lines indicate best fits for participants trained on lattice (purple) and modular (blue) graphs. (a) Interaction effect of graph × average recruitment on response time. (b) Interaction effect of graph limbic recruitment on response time. (c) Interaction effect of graph × average integration on response time. (d) Interaction effect of graph × limbic and temporoparietal integration on response time. (e) Interaction effect of graph × limbic and subcortical integration on response time. (f) Interaction effect of graph default mode and temporoparietal integration on response time. (g) Interaction effect of run × ventral attention and temporoparietal integration on response time. Each data point represents a participant; graded purple and blue lines indicate best fit for each task run.

A significant graph × integration effect at the global level also followed the same trend as the graph × recruitment effect; high average integration was associated with lower response times for participants in the modular group (*t* = −2.254, *df* = 120.3, *p*_FDR_ = 0.026; Fig. 6c). Three graph × integration effects were significant at the system level, all exhibiting similar trends as average integration: graph × limbic-temporoparietal (*t* = −4.488, *df* = 123.1, *p*_FDR_ = 0.00059; Fig. 6d), limbic-subcortical (*t* = −2.973, *df* = 120.23, *p*_FDR_ = 0.04272; Fig. 6e), and default mode-temporoparietal (*t* = −3.729, *df* = 123.1, *p*_FDR_ = 0.00525; Fig. 6f) integration. The fact that these graph × integration effects on response time show the same pattern as the recruitment models suggests that the association between high recruitment values and successful performance for the modular group does not indicate a segregation of cognitive systems from each other, but that interactions between intact cognitive systems subserved performance on this task for participants trained on the modular graph. Specifically, successful participants in the modular group tended to have higher integration of the limbic system with the temporoparietal and subcortical systems, as well as higher integration of the temporoparietal system with the default mode system.

We further investigated the relationship between response time and limbic, temporoparietal, default mode, and subcortical integration at the parcel level (Fig. 7). We found significant graph × limbic-temporoparietal integration interaction effects for the integration of three parcel pairs: the integration of a left hemisphere temporal pole parcel with a left hemisphere superior temporal gyrus parcel (LH LimbicA TempPole 2, LH TempPar 3; *t* = −4.812, *df* = 120.543, *p*_FDR_ = 0.00931), the integration of a left hemisphere orbital frontal cortex parcel with a right hemisphere superior temporal sulcus parcel (LH LimbicB OFC 4, RH TempPar 5; *t* = −3.698, *df* = 125.81, *p*_FDR_ = 0.0331), and the integration of a right hemisphere temporal pole parcel with a right hemisphere superior temporal sulcus parcel (RH LimbicA TempPole 1, RH TempPar 2; *t* = −3.534, *df* = 120.133, *p*_FDR_ = 0.0426). We found significant graph × limbic-subcortical integration interaction effects for the integration of 2 parcel pairs: the integration of a left hemisphere temporal pole parcel with a right hemisphere amygdala parcel (LH LimbicA TempPole 4, RH lAMY; *t* = −3.891, *df* = 121.064, *p*_FDR_ = 0.0251) and the integration of a right hemisphere temporal pole parcel with a left hemisphere hippocampus parcel (RH LimbicA TempPole 5, LH pHIP; *t* = −4.468, *df* = 119.384, *p*_FDR_ = 0.0127). While there were no significant effects of limbic-default mode integration on response time at the system level, parcel level analyses revealed significant graph × limbic-default mode integration interaction effects for the integration of 28 parcel pairs (−5.238 < *t* < −3.521, 119.301 < *df* < 121.16, 0.008 < *p*_FDR_ < 0.0434). All limbic parcels associated with this effect were located in the right hemisphere temporal pole, while default mode parcels were distributed across the left hemisphere default mode system. Together, these findings support the conclusion that the system-level graph × limbic-temporoparietal and limbic-subcortical integration effects on response time were likely driven by interactions of the temporal pole with the amygdala and superior temporal gyrus and sulcus, and indicate that the interaction of the temporal pole with the default mode system was also associated with lower response times for the modular group despite no system level effects.

**Figure 7.**
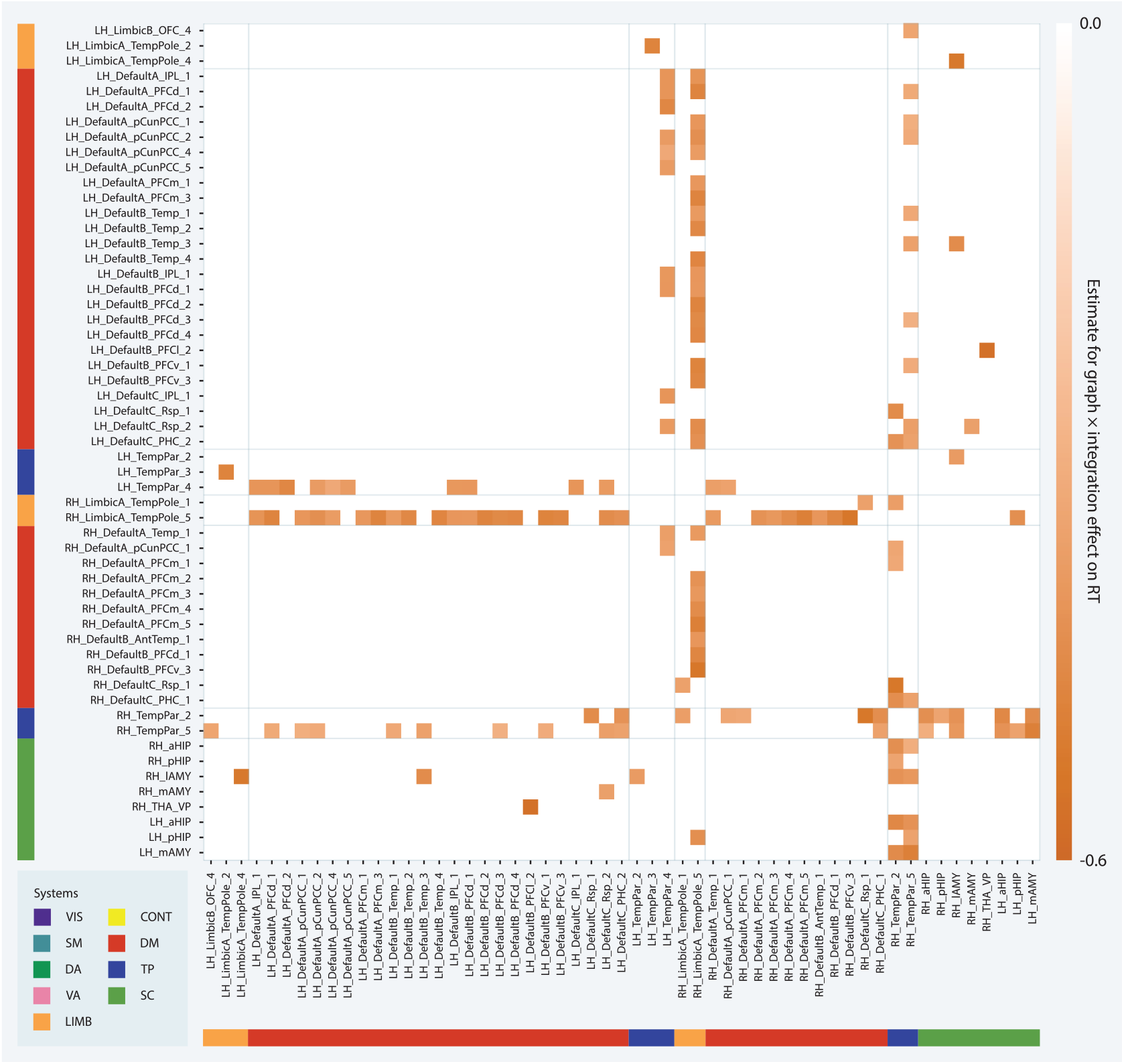
Parcel level network predictors of response time. (a–f) Interaction effect of graph × integration on response time for parcel level integration values. Each entry on the x- and y-axes corresponds to a parcel. Each square of the matrix indicates the estimate for the effect of graph × integration according to the corresponding multilevel model. The darker the square, the more negative the relationship between response time and integration for the modular group as compared to the lattice group.

We found significant graph × default mode-temporoparietal integration interaction effects for the integration of 28 parcel pairs (−5.26 < *t* < −3.532, 119.441 < *df* < 128.332, 0.008 < *p*_FDR_ < 0.0426). All temporoparietal parcels were located in the superior temporal sulcus, 78% of which were located in the posterior superior temporal sulcus, while default mode parcels were distributed across all components of the default mode system except the lateral prefrontal cortex. Most (75%) of these pairs included a left hemisphere default mode parcel, whereas superior temporal sulcus parcels were distributed across both hemispheres. These results suggest that the superior temporal sulcus and left hemisphere default mode system played a central role in the observed graph × default mode-temporoparietal integration effect on response time. No significant default mode-subcortical integration effects on response time were found at the system level, but at the parcel level, significant graph × integration effects were found for the integration of a left hemisphere superior temporal sulcus parcel with a right hemisphere amygdala parcel (DefaultB Temp 3, RH lAMY; *t* = −3.961, *df* = 119.931, *p*_FDR_ = 0.0221) and the integration of a left hemisphere retrosplenial cortex parcel with a right hemisphere amygdala parcel (DefaultC Rsp 2, RH mAMY; *t* = −3.614, *df* = 119.288, *p*_FDR_ = 0.0378). Similarly, no significant temporoparietal-subcortical integration effects on response time were found at the system level, but at the parcel level, significant graph × integration effects were found for the integration of two right hemisphere superior temporal sulcus parcels with an identical set of amygdala and hippocampus parcels (RH TempPar 2, RH TempPar 5, RH lAMY, LH mAMY, LH aHIP, RH aHIP, LH pHIP; −4.962 < *t* < −3.594, 120.319 < *df* < 122.682, 0.0075 < *p*_FDR_ < 0.0392), as well as the integration of a left hemisphere superior temporal sulcus parcel with a right hemisphere amygdala parcel (RH TempPar 2, RH lAMY; *t* = −3.696, *df* = 119.057, *p*_FDR_ = 0.0333). The overlap in regions showing significant graph × integration effects on response time in parcel level analyses of limbic, default mode, temporoparietal, and subcortical integration and the fact that these effects are all in the same direction provide evidence for a network of regions including the superior temporal sulcus, temporal pole, amygdala, hippocampus, and left hemisphere default mode system whose integration supported successful performance for the modular group.

We also found a significant interaction effect of task run × temporoparietal-ventral attention integration at the system level (*t* = 5.141, *df* = 118.18, *p*_FDR_ < 0.0001) indicating that the relationship between temporoparietal-ventral attention integration and response times became less negative over time (Fig. 6g).

When we investigated this relationship at the parcel level we also found significant run × ventral attention-temporoparietal integration interaction effects for the integration of six parcel pairs (3.508 < *t* < 4.449, 118.178 < *df* < 118.742, 0.0212 < *p*_FDR_ < 0.0495). All pairs included the same right hemisphere superior temporal sulcus parcel (RH TempPar 5). The ventral attention parcels included bilateral parcels encompassing parts of the midcingulate cortex, premotor cortex, and frontal eye fields (SalVentAttnA FrMed 1), a right hemisphere frontal operculum parcel (RH SalVentAttnA FrOper), a right hemisphere precentral gyrus parcel (RH SalVentAttnA PrC 1), and three right hemisphere insula parcels (RH SalVentAttnA Ins 3, RH SalVentAttnA Ins 2, RH SalVentAttnB Ins 2). Overall, these findings indicate that the integration of the superior temporal sulcus with a set of ventral attention parcels similar to those whose integration with the visual system increased over time tracked with individual differences in response time during early learning.

## 4 Discussion

The present work provides the first time-resolved network analysis of fMRI data collected during a graph learning task comparing graphs with differing topologies. We sought to provide a description of the observed networks in order to situate this study in the functional connectivity literature broadly, as well as to address the hypotheses that the functional brain networks that support the learning of probabilistic sequences generated via random walks on a graph (1) overlap with those implicated in other statistical learning studies and (2) depend on the topology of the underlying graph. In testing these hypotheses, we uncovered three key results. First, functional connectivity networks constructed from task data were more stable and integrated than networks constructed from resting state data. Second, the integration of visual and ventral attention regions increased over task runs as learning occurred. Third, an interaction of graph type and connectivity of limbic, default mode, temporoparietal, and subcortical systems predicted response times. In the remainder of this section, we will place these three findings within the context of the broader literature before outlining important methodological considerations and directions for future work.

### 4.1 Task networks were more stable and integrated than rest networks

In comparison to functional networks derived from resting-state scans, task networks exhibited greater stability, widespread changes in recruitment, and higher overall integration. Consistent with previous task-based functional connectivity analyses, static functional connectivity was higher across all systems during task performance [59]. Multilayer community detection in time-resolved networks revealed a general decrease in flexibility in task networks driven by stable connectivity patterns in the dorsal attention, limbic, frontoparietal control, and default mode systems. Prior work has also found such increases in the stability of task networks, which has been hypothesized to reflect increases in cognitive constraints [59, 60, 61]. The increase in stability of the higher-order and hetero-modal systems contrasted with the increase in flexibility of the visual system in task networks. Recruitment of the visual system also decreased dramatically during task performance while the integration of the visual system with all other systems increased; these findings together indicate that the visual system is a flexible integrator during visuomotor graph learning. Decreases in within-network connectivity of the visual system and increases in between-network connections involving the visual system have been reported previously for visual task performance [59, 62, 63]; the relative flexibility of the visual system appears to be specific to the graph learning task [64]. Recruitment of the dorsal attention, limbic, default mode, and subcortical systems was higher during task performance, reflecting a higher degree of cohesiveness. However, all nine systems exhibited increases in inter-system integration with at least three other systems, suggesting that task performance required a high degree of inter-system coordination. The widespread increase in integration across systems we observed during task has been reported for many other tasks [59, 65]. We also observed selective decreases in integration of the default mode system with the frontoparietal control and dorsal attention systems. As the frontoparietal control and dorsal attention systems are associated with goal-directed cognition, this effect could be interpreted as the segregation of a task-negative system [66]. However, as discussed below, an alternative interpretation regards the frontoparietal control system as less integral to this implicit learning task and the default mode system as critical for representing the graph structure underlying stimulus presentation, especially for participants learning the modular graph.

### 4.2 Integration of visual and ventral attention regions increased over task runs

Previous task-based functional connectivity work has suggested that task-relevant networks tend to be more segregated during late training than early training, and this change has been associated with an increase in the automaticity of task performance [51, 53]. Short deterministic sequence production tasks such as that employed in [51] allow for fully autonomous motor production as a single visual cue determines all of the proceeding motor responses. This is evidenced by the functional segregation of the visual and motor systems observed in [51]. The graph learning task described here provides a statistically structured sequence such that motor prediction and preparation should be possible, but we expected that visual regions would never be able to fully disengage from motor regions. While we did not observe any system-level effects of task run on recruitment or integration, 157 pairs of parcels showed changes in integration over time, 82% of which became more integrated and 18% of which became less integrated. Of the parcel pairs that became more integrated, most included a visual system parcel, with visual-ventral attention and visual-somatomotor integration accounting for the majority of such pairs. Most of the parcel pairs that became less integrated over time were composed of a parcel in the frontoparietal control system and either a visual or dorsal attention parcel. These results suggest that (1) efficient performance of a graph learning task may require the relevant systems to remain integrated, even during late learning, and (2) automaticity, in this case, may manifest as a switch from the top-down influence of cognitive control networks on the visual system to bottom-up influence of the stimulus-driven ventral attention system. The ventral attention system also includes regions in the inferior frontal gyrus and insula, both of which have been shown to exhibit similar neural representations for stimuli within the same community in a modular graph [35]. Furthermore, inferior frontal gyrus activity measured by intracranial EEG has previously been associated with representations of the latent space in a graph learning task similar to ours [48]. Statistical coupling of stimuli should arise after multiple exposures, which could explain why integration between the visual and ventral attention regions increased over run. Consistent with this hypothesis, the inferior frontal gyrus and insula show a repetition enhancement effect, wherein neural activity increases as more time is spent within a single community [35].

### 4.3 Interaction of graph type and connectivity of limbic, default mode, temporoparietal, and subcortical systems predicts response times

We found differences between the two graph groups with respect to the relationship between response times and recruitment of the limbic system, integration of the limbic system with the temporoparietal system and subcortex, and integration of the default mode system with the temporoparietal system. Higher recruitment and integration values for these systems were associated with lower response times for the modular group and higher response times for the lattice group, consistent with our hypothesis that distinct brain networks support graph learning depending on the topology of the underlying graph. Prior graph learning work has shown that functional connectivity within the limbic system is higher during trials which represent a transition from one module to another for participants training on a modular graph [47]. Our finding that recruitment of the limbic system is associated with lower response times for the modular group extends this result by showing that connectivity within the limbic system has behavioral relevance. Individual regions within the limbic, temporoparietal, default mode, and subcortical systems have also previously been implicated in statistical learning. As mentioned above, neural representations of statistically coupled items are markedly similar in the inferior frontal gyrus, which is also partially contained within the default mode system [35]. Similar neural representations of statistically coupled items have also been found within the superior temporal gyrus, encompassed by the limbic, default mode, and temporoparietal systems (in addition to the somatomotor system), the anterior temporal lobe, which is categorized as limbic, and the hippocampus, which is a subcortical structure [35]. The superior temporal and inferior frontal gyri have also been associated with learning statistically structured sequences as compared to statistically unstructured sequences [10]. Thus, the relationship between integration of the limbic, default mode, temporoparietal, and subcortical systems and response times may reflect their involvement in representing the topology of the graphs underlying sequence generation, particularly for participants in the modular group.

Investigating the interaction effect of graph and integration of the limbic, default mode, temporoparietal, and subcortical systems on response time at the parcel level indicated that the integration of a network of regions including the superior temporal sulcus, temporal pole, amygdala, hippocampus, and left hemisphere default mode system may have supported successful performance for the modular group. The temporal pole has been implicated in the processing and memory of a variety of complex stimuli, including complex objects [67]. There is also evidence that the temporal pole encodes information regarding stimulus novelty and the entropy of a sequence [68, 69]. The amygdala, which is in close anatomical proximity to the temporal pole, is thought to assign uncertainty values to representations [70, 71]. Relevantly, we found a graph × integration effect on response time for integration of amygdala and temporal pole parcels that showed that integration of these regions was associated with lower response times for the modular group but not the lattice group, perhaps relating to the difference in statistical information contained in the structure of each graph. The finding that amygdala activity was also strongly associated with representations of the latent space in another aforementioned graph learning task is compelling support for this hypothesis [48].

Recent functional and effective connectivity studies have implicated the superior temporal sulcus in a ventrolateral visual stream for invariant object recognition [72, 73]. These studies further suggest that the superior temporal sulcus receives inputs from reward-related ventromedial prefrontal cortex regions and has effective connectivity with memory-related posterior cingulate cortex regions, both of which have regions in the default mode system. Integration of parcels in these regions with superior temporal sulcus parcels showed significant graph × integration effects on response time, indicating that the connectivity of this network may be important for learning on modular graphs. Posterior cingulate and adjacent retrosplenial cortical regions are thought to play a role in spatial navigation [72] and, along with medial prefrontal cortex and temporoparietal regions, have been associated with the representation of conceptual spaces [74]. Lesion and fMRI studies have also identified the medial temporal lobe as a key, and perhaps necessary region for statistical learning, and graph learning in particular [45, 32, 31, 26]. The hippocampus, which is in the medial temporal lobe, is more active when traversing edges between nodes within the same community in a modular graph and less active at community boundaries [45]. Additionally, neural representations of items in the same community are markedly similar in the hippocampus. Together, these observations suggest that the hippocampus may be particularly important for representing community structure. Moreover, the hippocampus has long been implicated in the representation of cognitive maps [75]. Given the role of the hippocampus in representing spatial and other relational information, as in the cases of episodic memory and spatial navigation, the cytoarchitecture of the region is plausibly capable of storing graphs generated by temporal statistics [37, 76, 77, 78]. Supporting this hypothesis, [46] found that patterns of functional connectivity between the medial temporal lobe, postcentral gyrus, and lateral occipital cortex during graph learning are associated with the accuracy with which trial identity can be decoded in the hippocampus and postcentral gyrus. As graph learning can be construed as a process involving the exploration of conceptual space, we posit that successful learners in the modular group recruited a network connecting areas needed for object recognition with areas important for spatial navigation to a higher degree than participants in the lattice group.

### 4.4 Methodological considerations

Certain methodological considerations are relevant when interpreting the results discussed above. Here, we focused on just two graph types: modular and ring lattice. Comparing response time and functional connectivity patterns reported here with those associated with other types of graphs is a compelling direction for future studies because it could help elucidate which topological properties are optimal for learning. Future work would also benefit from exploring how connectivity patterns might differ according to how a graph is traversed; although all sequences were generated via random walks in the dataset employed here, variations in how a graph is traversed have been shown to impact the learnability of a graph [41, 79]. Furthermore, the learning period we analyzed here was approximately 30 minutes. By examining the relation between behavior and connectivity patterns over a learning period lasting multiple days or weeks, future work could clarify how this relation changes over the different stages of learning and how it is impacted by the process of consolidation [80, 81, 82, 51, 83]. Finally, although we related the observed connectivity patterns to observed behavioral responses, we did not examine how task structure may have influenced the amount of shared information between brain regions. A psychophysiological interaction analysis would complement the results presented here by taking into account the experimental context in which the interactions between brain regions occurred [84].

### 4.5 Conclusion

This study leverages time-resolved network analysis techniques to investigate the connectivity and community structure of large-scale cognitive systems during visuomotor graph learning. In examining the relationship between task performance and community structure for participants learning on graphs with different topologies, we found that integration between the limbic, default mode, temporoparietal, and subcortical systems, as well as recruitment of the limbic system, was associated with lower response times for participants learning on the modular graph but not the lattice graph. These results expand the literature elucidating the neural correlates of previously reported differences in behavior, providing a novel, time-resolved connectivity-based perspective [46, 43, 45, 35]. Broadly, this work contributes to knowledge of how humans process complex statistical structures and paves the way for future work addressing other graph topologies.

## 5 Methods

### 5.1 Dataset

The dataset used here was first reported by Kahn et al. [46]. Here we provide a brief summary and refer the reader to that report for more details.

#### 5.1.1 Participants

Thirty-four right-handed individuals (aged 18–34 years, 68% female) were recruited from the general population of Philadelphia, Pennsylvania. All participants provided written informed consent and all procedures were approved by the Institutional Review Board at the University of Pennsylvania according to the Declaration of Helsinki. Data from thirty participants were included in our final analysis. Four individuals were excluded due to (1) failure to complete pre-training (see Section 5.1.4), (2) task performance at chance levels, (3) technical difficulties that prevented the recording of behavioral data, or (4) failure to complete all fMRI scanning sessions.

#### 5.1.2 Imaging

##### Acquisition

Each participant underwent seven BOLD fMRI scans, acquired with a 3T Siemens Magnetom Prisma scanner using a 32-channel head coil. Two pre-task scans were acquired as participants rested inside the scanner, followed by five task scans and two post-task resting-state scans. Imaging parameters were based on the ABCD protocol [85]. Each scan employed a 60-slice gradient-echo echo-planar imaging sequence with 2.4 × 2.4 × 2.4 mm isotropic voxels, an iPAT slice acceleration factor of 6, and an anterior-to-posterior phase encoding direction; the repetition time (TR) was 800 ms and the echo time (TE) was 30 ms. A T1w reference image was acquired using an `MEMPRAGE`sequence [86] with the following parameters: 2530 ms TRs, 1.69 ms, 3.55 ms, 5.41 ms, and 7.27 ms TEs, 1 × 1 × 1 mm voxels, anterior-to-posterior encoding, and a GRAPPA acceleration factor of 3 [87].

##### Preprocessing

Results included in this manuscript come from preprocessing performed using *fMRIPrep* 20.1.0 [88, 89], which is based on *Nipype* 1.4.2 [90, 91].

The T1-weighted (T1w) image was corrected for intensity non-uniformity (INU) with `N4BiasFieldCorrection`[92], distributed with ANTs 2.2.0 [93], and used as the T1w-reference throughout the workflow. The T1w-reference was then skull-stripped with a *Nipype* implementation of the `antsBrainExtraction.sh`workflow (from ANTs), using OASIS30ANTs as the target template. Brain tissue segmentation of cerebrospinal fluid (CSF), white-matter (WM), and gray-matter (GM) was performed on the brain-extracted T1w using `fast`[94]. Brain surfaces were reconstructed using `recon-all`[95], and the brain mask estimated previously was refined with a custom variation of the method to reconcile ANTs-derived and FreeSurfer-derived segmentations of the cortical gray-matter of Mindboggle [96]. Volume-based spatial normalization to two standard spaces (MNI152NLin2009cAsym, MNI152NLin6Asym) was performed through nonlinear registration with `antsRegistration`(ANTs 2.2.0), using brain-extracted versions of both the T1w reference and the T1w template. The following templates were selected for spatial normalization: *ICBM 152 Nonlinear Asymmetrical template version 2009c* [97] and *FSL’s MNI ICBM 152 non-linear 6th Generation Asymmetric Average Brain Stereotaxic Registration Model* [98].

For each of the 7 BOLD runs per participant, the following preprocessing was performed. First, a reference volume and its skull-stripped version were generated using a custom methodology of *fMRIPrep*. Head-motion parameters with respect to the BOLD reference (transformation matrices, and six corresponding rotation and translation parameters) were estimated before any spatiotemporal filtering using `mcflirt`[99]. BOLD runs were slice-time corrected using `3dTshift`from AFNI 20160207 [100]. A B0-nonuniformity map (or *fieldmap*) was estimated based on two (or more) echo-planar imaging (EPI) references with opposing phaseencoding directions, with `3dQwarp`[100] (AFNI 20160207). Based on the estimated susceptibility distortion, a corrected EPI (echo-planar imaging) reference was calculated for a more accurate co-registration with the anatomical reference. The BOLD reference was then co-registered to the T1w reference using `bbregister`(FreeSurfer), which implements boundary-based registration [101]. Co-registration was configured with nine degrees of freedom to account for distortions remaining in the BOLD reference.

The BOLD time series (including slice-timing correction when applied) were resampled onto their original, native space by applying a single composite transform to correct for head motion and susceptibility distortions. We refer to these resampled BOLD time-series as the *preprocessed BOLD in original space*, or just *preprocessed BOLD*. The BOLD time-series were resampled into several standard spaces, correspondingly generating the following *spatially-normalized, preprocessed BOLD runs*: MNI152NLin2009cAsym, MNI152NLin6Asym. A reference volume and its skull-stripped version were generated using a custom methodology of *fMRIPrep*.

Several confounding time series were calculated based on the *preprocessed BOLD* : framewise displacement (FD), DVARS, and three region-wise global signals. FD was computed using two formulations: absolute sum of relative motions [102] and relative root mean square displacement between affines [99]. FD and DVARS were calculated for each functional run, both using their implementations in *Nipype* [102]. The three global signals were extracted within the CSF, WM, and whole-brain masks. Additionally, a set of physiological regressors were extracted to allow for component-based noise correction [103].

Principal components were estimated after high-pass filtering the *preprocessed BOLD* time-series (using a discrete cosine filter with 128s cut-off) for the two *CompCor* variants: temporal (tCompCor) and anatomical (aCompCor). tCompCor components were then calculated from the top 5% variable voxels within a mask covering the subcortical regions. This subcortical mask was obtained by heavily eroding the brain mask, which ensures it did not include cortical GM regions. For aCompCor, components were calculated within the intersection of the aforementioned mask, and the union of CSF and WM masks was calculated in T1w space after their projection to the native space of each functional run (using the inverse BOLD-to-T1w transformation). Components were also calculated separately within the WM and CSF masks. For each CompCor decomposition, the *k* components with the largest singular values were retained, such that the retained components’ time series were sufficient to explain 50 percent of variance across the nuisance mask (CSF, WM, combined, or temporal). The remaining components were removed from consideration. The head-motion estimates calculated in the correction step were also placed within the corresponding confounds file. The confound time series derived from head motion estimates and global signals were expanded with the inclusion of temporal derivatives and quadratic terms for each [104]. Frames that exceeded a threshold of 0.5 mm FD or 1.5 standardized DVARS were annotated as motion outliers. All resamplings were performed with *a single interpolation step* by composing all the pertinent transformations (i.e., head-motion transform matrices, susceptibility distortion correction when available, and co-registrations to anatomical and output spaces). Gridded (volumetric) resamplings were performed using `antsApplyTransforms`(ANTs), configured with Lanczos interpolation to minimize the smoothing effects of other kernels [105]. Non-gridded (surface) resamplings were performed using `mri vol2surf`(FreeSurfer).

Many internal operations of *fMRIPrep* used *Nilearn* 0.6.2 [106], mostly within the functional processing workflow. For more details of the pipeline, see the section corresponding to workflows in *fMRIPrep*’s documentation.

#### 5.1.3 Stimuli

Visual stimuli were displayed on either a laptop screen (see Section 5.1.4) or on a screen inside the MRI machine (see Sections 5.1.5) and 5.1.6) and consisted of fifteen unique abstract shapes generated by perturbing a sphere with sinusoids using the `MATLAB`package `ShapeToolbox`(Fig. 1a, *left*). Five variations of each shape were also created through scaling and rotating, and the appearance of variations was balanced within task runs so as to capture neural activity invariant to these properties. Each shape was paired with a unique motor response, with possible motor responses spanning the set of all fifteen one- or two-button chords on a five-button response pad (Fig. 1a, *right*). The five buttons on the response pad corresponded to the five fingers on the right hand: the leftmost button corresponded to the thumb, the second button from the left corresponded to the index finger, and so on. In addition to the shape cue, motor responses were explicitly cued by a row of five square outlines on the screen, each of which corresponded to a button on the response pad. Squares corresponding to the cued buttons turned red; for instance, if the first and fourth squares turned red, a correct response consisted of the thumb and ring finger buttons being pressed.

Participants were assigned one of two graph types: modular or ring lattice, both of which were composed of fifteen nodes and thirty undirected edges such that each node had exactly four neighbors (Fig. 1b, *left*). Nodes in the modular graph were organized into three densely connected five-node clusters, and a single edge connected each pair of clusters. Each node in a participant’s assigned graph was associated with a stimulus-response pair (Fig. 1b, *right*). While the stimulus and response sets were the same for all participants, the stimulus-response-node mappings were random and unique to each participant. This mapping was consistent across runs for an individual participant. The order in which stimuli were presented during a single task run was dictated by a random walk on a participant’s assigned graph; that is, an edge in the graph represented a valid transition between stimuli. No other meanings were ascribed to edges. Walks were randomized across participants as well as across runs for a single participant. Two constraints were instituted on the random walks: (1) all walks were required to visit each node at least ten times, and (2) walks on the modular graph were required to include at least twenty cross-cluster transitions.

#### 5.1.4 Pre-training

Prior to entering the scanner, participants performed a “pre-training” session during which stimuli were displayed on a laptop screen and motor responses were logged on a keyboard using the keys ‘space’, ‘j’, ‘k’, ‘l’, and ‘;’. Participants were provided with instructions for the task, after which they completed a brief quiz to verify comprehension. To facilitate the pre-training process, shape stimuli were divided into five groups of three. First, participants were simultaneously shown a stimulus and its associated motor response cue, and then were asked to press the associated button(s) on the keyboard. Next, participants were shown the stimulus alone and asked to perform the motor response from memory. If they did not respond within two seconds, the trial was repeated. If they responded incorrectly, the motor response cue was displayed. Once the participant responded correctly, the same procedure was repeated for the other two stimuli in the subset. Then, participants were instructed to respond to a sequence of the same three stimuli, presented without response cues. The sequence was repeated until the participant responded correctly for 12 consecutive trials. This entire process was repeated for each three-stimulus subset. The three stimuli in a given subset were always chosen so that none were adjacent to one another in the graph dictating the order of the sequence used during the “Training” stage (see below).

#### 5.1.5 Training

Five runs of 300 trials each were conducted inside the MRI scanner. During each trial, participants were presented with a visual stimulus consisting of a row of five square outlines, each of which contained the same abstract shape (Fig. 1c.). Five hundred milliseconds later, one or two of the squares turned red. Using a five-button controller, participants were required to press the button(s) corresponding to the red square(s) on the screen. If a participant pressed the wrong button(s), the word “Incorrect” appeared on the screen. The following stimulus did not appear until the correct buttons were pressed. Participants were instructed to respond as quickly as possible. Sequences were generated via random walks on the participants’ assigned graphs.

#### 5.1.6 Recall

This task was originally designed to test hypotheses regarding how the different graph structures used to generate trial sequences influenced participants’ neural representations of the stimuli [46]. To allow the experimenters to separate representations of the motor responses from representations of the visual stimuli and to assess learning, participants completed additional “recall” trials during which shapes were presented without their corresponding motor response cues. Since the hypotheses addressed in this manuscript focus on the neural correlates of the learning process, the only runs analyzed were the training runs described above.

At the end of each training run, participants completed 15 additional “recall” trials during which each shape was presented without a motor response cue, requiring participants to recall the correct response. Participants were given two seconds to respond; if they did not respond within this time, the trial was marked as incorrect. If they responded incorrectly, the trial was marked as incorrect and the motor response cue was displayed below the shape for the remainder of the two seconds. If they responded correctly, the outline of the square containing the shape turned green. A six-second delay separated the end of the last recall trial in a run from the start of the first training trial in the subsequent run. During a second scanning session (not described in the “Imaging” section above), participants completed another eight task runs, each comprised of 60 recall trials. The orderings of all recall trials were also determined by walks on the participants’ assigned graphs, although the walks were Hamiltonian instead of random for 120 of the recall trials conducted during the second session.

### 5.2 Network analysis

#### 5.2.1 Network construction

##### Parcellation

We employed three well-established atlases to parcellate the functional imaging data into regions of interest (ROIs, Fig. 2b). Consistent with recent task-based fMRI studies [107, 53], we used the 300-parcel Schaefer parcellation to assign non-overlapping cortical ROIs to 17 large-scale functional systems: two visual, two somatomotor, two dorsal attention, two ventral attention, two limbic, three frontoparietal control, three default mode, and one temporoparietal [54]. For completeness, and because learning, memory, and motor networks were likely to be implicated in task execution [108, 109, 110], we also employed subcortical and cerebellar atlases. For ROIs in the hippocampus, thalamus, amygdala, caudate nucleus, nucleus accumbens, putamen, and globus pallidus, we used the Scale II, 3T version of the 32-parcel Melbourne atlas [56]. To parcellate the cerebellum, we chose the 17-parcel Buckner atlas, which assigns a cerebellar ROI to each of the 17 cortical systems implemented in the Schaefer cortical atlas [55]. For each of the 349 total ROIs, the BOLD signal from the subsumed voxels was extracted and averaged using FSL’s `fslmeants`function to obtain one time series per region. When examining system-level effects, we collapsed the 17 cortical/cerebellar systems into eight systems by merging systems with the same functional role; that is, by merging the two visual systems, the two somatomotor systems, the two dorsal attention systems, the two ventral attention systems, the three frontoparietal control systems, and the three default mode systems [107]. In these system-level analyses, subcortical parcels composed a single ninth system.

##### Static network construction

A functional connection is the statistical relationship between time series of neurophysiological activity for a given pair of brain regions [111]. While this statistical relationship has been defined by several different measures in the literature, the product-moment correlation coefficient (Pearson *r*) is the most common [112, 113]. Compared to other statistical association metrics, the product-moment correlation coefficient is also more sensitive to modular structure in functional networks, which was a key focus of our study [112]; see Section 5.2.2. By defining nodes as ROIs and edges between nodes as the product-moment correlation coefficient for the time series of the connected nodes, we constructed a 349 × 349 functional connectivity adjacency matrix for each fMRI scan. It has been shown that setting negative product-moment values to zero improves the reliability of the measure for functional connectivity analyses as negative product-moment values are more sensitive to motion artifacts; thus, we set any negative product-moment values in each functional connectivity matrix to zero.

##### Time-resolved network construction

To test our hypotheses concerning the relationship between functional connectivity patterns and learning, we constructed time-resolved multilayer networks (Fig. 1c, d). For each time series, we obtained four multilayer networks generated using a traditional sliding-window approach [114, 115, 50, 116], which differed only in the degree of overlap between adjacent time windows. The aim of varying the degree of overlap was to increase the rigor of our method and enhance the reliability of our conclusions, which we drew only when findings were replicated across network construction methods.

We parsed each time series into multiple overlapping time windows and calculated 349 × 349 functional connectivity matrices for each window using product-moment correlations as described above, yielding multilayer networks wherein a single layer corresponded to a single time window. To ensure that each window would capture biologically relevant signal fluctuations, we adhered to an established guideline suggesting that windows be at least as long as the inverse of the lowest frequency sampled [117]. As we applied a bandpass filter of 0.01–0.08 Hz, we thus obtained windows that were 100 s in length, corresponding to 125 TRs. The overlap between adjacent windows was set to 1, 32, 63, or 124 TRs; a new network was generated for each setting, resulting in four sliding-window multilayer networks for each time series.

For simplicity, the time-resolved network statistics presented in the main text are associated solely with networks generated using an overlap of 124 TRs [118]. However, we only report and discuss results that are replicated in all four varieties of multilayer networks.

#### 5.2.2 Network diagnostics

#### Community detection

Given the relevance of modularity for cognition [119, 50, 120], we examined the modular architecture of the time-resolved multilayer networks (Fig. 1e). In a fully connected network, modules are groups of nodes that have stronger within-group connections than between-group connections. By analyzing the modular properties of the time-resolved networks, we estimated which brain regions preferentially modified their interactions as a function of task execution. To identify such modules, we applied a generalized Louvain community detection algorithm [53, 51, 57, 50, 121]. This algorithm operates by maximizing a common multilayer modularity quality function, defined as follows:

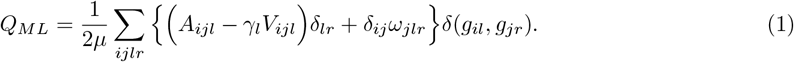

Here *A*_*ijl*_ is the product-moment correlation between nodes *i* and *j* in layer *l. V_ijl_* is the associated element of the Newman-Girvan null model 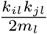 in which the total edge weight in layer *l* is 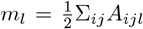, *k*_*il*_ is the intra-layer strength of node *i* in layer *l*, and *k*_*jl*_ is the intra-layer strength of node *j* in layer *l. γ_l_* is the structural resolution parameter of layer *l*, which influences the number of communities detected, and *ω*_*jlr*_ is the strength of the link between node *j* in layer *r* and node *j* in layer *l*, which affects the temporal stability of the detected community structure. The total edge weight is 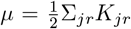, where *K*_*jl*_ = *κ*_*jl*_ + *c*_*jl*_ is the strength of node *j* in layer *l*, and *c*_*jl*_ =Σ_*r*_*ω*_*jlr*_ is the inter-layer strength of node *j* in layer *l*. The quantity *g*_*il*_ is the community assignment of node *i* in layer *l*, and *g*_*jr*_ is the community assignment of node *j* in layer *r*. Both *γ*_*l*_ and *ω*_*ijl*_ were set to 1, in accordance with similar analyses of time-resolved functional connectivity [53, 52, 50]. As each individual run of this algorithm produces a slightly different result with multiple partitions reaching near-optimal values of *Q* [122], we ran the algorithm 100 times, obtaining 100 different partitions of brain regions into communities (or modules) for each layer of the time-resolved networks. The statistics below were calculated for each of the 100 partitions and then averaged across partitions [50].

##### Flexibility

To determine the degree to which the communities identified by the community detection algorithm were stable across time windows, we calculated the flexibility [50], *f*, of node *i* as follows:

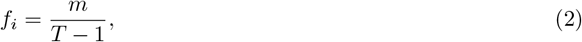

where *m* is the number of times node *i* changed communities, and *T* is the number of time windows in a given multilayer network [123]. A high flexibility value suggests that the interactions exhibited by the node in question were temporally variable or fluid.

##### Module allegiance

To summarize the time-resolved modular structure identified by the community detection algorithm, we constructed a module allegiance matrix [61, 51, 50], *P*, for each community partition. Each element *P*_*ij*_ of a module allegiance matrix provides the frequency with which brain regions were assigned to the same communities across time windows. Mathematically, we can write the element of *P* as follows:

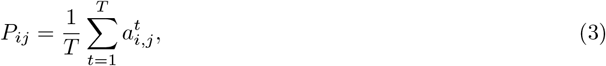

where *P*_*ij*_ is the fraction of network layers for which node *i* and node *j* were assigned to the same community and *T* is the number of layers. For each layer *t*,

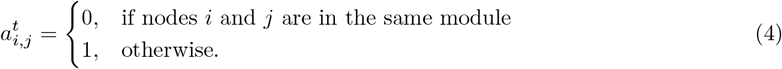

Module allegiance, as defined in this way, is hence a useful quantity that allows us to assess the extent to which different regions interacted with one another over time without relying on a particular community partition. From the module allegiance matrices, we calculated recruitment and integration (see below).

##### Recruitment

To assess the degree to which nodes belonging to the same system were assigned to the same communities, we calculated the recruitment [51, 64], *R*, of node *i* with respect to system *S*, as defined below:

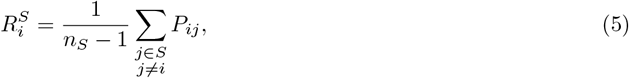

where node *i* and node *j* belong to system *S, P_ij_* is the fraction of network layers for which node *i* and node *j* belong to the same community and *n*_*S*_ is the number of nodes in *S*. When recruitment values are averaged across all nodes in a given system, a high value of *R*^*S*^ indicates strong within-system connections for system *S*, and a low value of *R*^*S*^ indicates weak within-system connections for system *S*. Recruitment can also be thought of as a measure of the cohesiveness or stability of a system [51]; if the nodes in a system are more strongly connected to each other than to nodes in other systems (see *Integration* below), then the system may be operating largely on its own and less in conjunction with other systems. When recruitment values are averaged across all nodes in a network, *R* gives the degree to which the communities identified by the community detection algorithm resemble the systems defined *a priori* by the parcellations described above.

##### Integration

To estimate inter-system interactions, we computed the integration [51, 64, 52], *I*, of system *S*_*k*_with system *S*_*l*_ as follows:

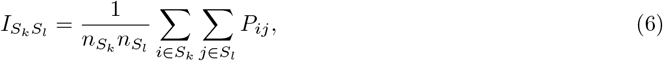

where *l* ≠ *k, P_ij_* is the fraction of network layers for which node *i* and node *j* belong to the same community, and 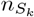 and 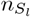 are the number of nodes in systems *k* and *l*, respectively. When integration values are averaged across all node pairs for a given pair of systems, a high value of integration is indicative of strong between-system connections. System level integration and recruitment are related in the following manner: if 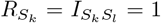, then all nodes in both *S*_*k*_ and *S*_*l*_ were always assigned to the same community, and so 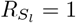, indicating both strong within-system and between-system connections for *S*_*k*_ and *S*_*l*_; if 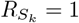 and 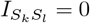, then all nodes in *S*_*k*_ were always assigned to the same individual community, to which nodes in *S*_*l*_ were never assigned; if 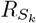 is low and 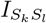 is high, then nodes in *S*_*k*_ were distributed across communities and nodes in *S*_*l*_ were distributed across the same communities (note that if 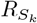 is low, then 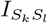 cannot be as high as if 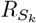 were high, because both *R* and *I* are higher when the relevant nodes are clustered in the same few communities than when they are distributed across many communities).

#### 5.2.3 Statistical modeling

We employed multilevel modeling in the following two contexts during our analyses. All models were implemented using the `R`package `lmerTest`[124].

##### Changes in brain network structure

To explore how brain network structure evolved over the course of the five task runs and differed between the two graph groups, we generated models of static functional connectivity, flexibility, recruitment, and integration with fixed effects of task run (one through five), graph type (modular or ring lattice), and the interaction between the two (graph × run). To account for individual variability, we added one random effect of participant to each model. The response variables, static functional connectivity, flexibility, recruitment, and integration, were defined on the global, system, and parcel levels: one model per response variable for global level results, one model per system (or pair of systems) per response variable for system-level results, and one model per parcel (or pair of parcels) per response variable for parcel level results. Estimated *p*-values were obtained via the Satterthwaite approximation, as implemented in the `lmerTest`package. For systemand parcel-level models, *p*-values obtained from models of the same network metric were corrected for multiple comparisons using the false-discovery rate procedure [125].

##### Relationship between network structure and behavior

To explore the relationships between network structure, graph type, and behavior over time, we generated models of response time with fixed effects of task run, graph type, and either static functional connectivity, flexibility, recruitment, or integration. For each network metric, we generated global-, system-, and parcel-level models with fixed effects of task run, graph type, network metric, and all the two- and three-way interactions between these. Each of the models also included one random effect of participant. Prior to model fitting, response time data were filtered to remove trials wherein (1) the response time was less than 100 ms, (2) the response time was more than 5 seconds, (3) the response time was more than 3 SDs away from that subject’s mean, and (4) the response was incorrect, as is consistent with previous work [46, 43, 40]. Estimated *p*-values were obtained via the Satterthwaite approximation, as implemented in the `lmerTest`package. Within each scale (global, system, or parcel) and network metric, *p*-values were corrected for multiple comparisons using the false-discovery rate procedure [125].

## 6 Data and code availability statement

Data and code used in this manuscript will be made available upon request.

## 7 Citation diversity statement

Recent work in several fields of science has identified a bias in citation practices such that papers from women and other minority scholars are under-cited relative to the number of such papers in the field [126, 127, 128, 129, 130, 131, 132, 133, 134]. Here we sought to proactively consider choosing references that reflect the diversity of the field in thought, form of contribution, gender, race, ethnicity, and other factors. First, we obtained the predicted gender of the first and last author of each reference by using databases that store the probability of a first name being carried by a woman [130, 135]. By this measure (and excluding self-citations to the first and last authors of our current paper), our references contain 14.29% woman(first)/woman(last), 9.18% man/woman, 20.75% woman/man, and 55.78% man/man. This method is limited in that a) names, pronouns, and social media profiles used to construct the databases may not, in every case, be indicative of gender identity and b) it cannot account for intersex, non-binary, or transgender people. Second, we obtained predicted racial/ethnic category of the first and last author of each reference by databases that store the probability of a first and last name being carried by an author of color [136, 137]. By this measure (and excluding self-citations), our references contain 8.9% author of color (first)/author of color(last), 18.99% white author/author of color, 16.81% author of color/white author, and 55.30% white author/white author. This method is limited in that a) names and Florida Voter Data to make the predictions may not be indicative of racial/ethnic identity, and b) it cannot account for Indigenous and mixed-race authors, or those who may face differential biases due to the ambiguous racialization or ethnicization of their names. We look forward to future work that could help us to better understand how to support equitable practices in science.

## 8 Acknowledgements

This research was funded by the National Institute of Health (Grant No. R21-MH-106799) and further supported by the Center for Curiosity. The content is solely the responsibility of the authors and does not necessarily represent the official views of any of the funding agencies.

## References

[1] Michael S. Vitevitch and Paul A. Luce. “Probabilistic Phonotactics and Neighborhood Activation in Spoken Word Recognition”. In: Journal of Memory and Language 40.3 (Apr. 1999), pp. 374–408. issn: 0749596X. doi: 10.1006/jmla.1998.2618. url: https://linkinghub.elsevier.com/retrieve/pii/S0749596×98926183.

[2] Alison Gopnik and Henry M. Wellman. “Reconstructing constructivism: Causal models, Bayesian learning mechanisms, and the theory theory”. In: Psychological Bulletin 138.6 (2012), pp. 1085–1108. issn: 00332909. doi: 10.1037/a0028044.

[3] Anna C. Schapiro and Nicholas B. Turk-Browne. “Statistical Learning”. In: Brain Mapping. Vol. 3. Elsevier, 2015, pp. 501–506. isbn: 9780123970251. doi: 10.1016/B978-0-12-397025-1.00276-1. url: https://linkinghub.elsevier.com/retrieve/pii/B9780123970251002761.

[4] József Fiser and Richard N. Aslin. “Statistical learning of higher-order temporal structure from visual shape sequences.” In: Journal of Experimental Psychology: Learning, Memory, and Cognition 28.3 (2002), pp. 458–467. issn: 1939-1285. doi: 10.1037/0278-7393.28.3.458. url: http://doi.apa.org/getdoi.cfm?doi=10.1037/0278-7393.28.3.458.

[5] Timothy F. Brady and Aude Oliva. “Statistical Learning Using Real-World Scenes”. In: Psychological Science 19.7 (2008), pp. 678–685. issn: 0956-7976. doi: 10.1111/j.1467-9280.2008.02142.x.

[6] Christopher M. Conway and Morten H. Christiansen. “Modality-constrained statistical learning of tactile, visual, and auditory sequences”. In: Journal of Experimental Psychology: Learning Memory and Cognition 31.1 (2005), pp. 24–39. issn: 02787393. doi: 10.1037/0278-7393.31.1.24.

[7] Jenny R. Saffran et al. “Statistical learning of tone sequences by human infants and adults”. In: Cognition 70.1 (1999), pp. 27–52. issn: 00100277. doi: 10.1016/S0010-0277(98)00075-4.

[8] Jenny R. Saffran, Richard N. Aslin, and Elissa L. Newport. “Statistical Learning by 8-Month-Old Infants”. In: Science 274.5294 (Dec. 1996), pp. 1926–1928. issn: 0036-8075. doi: 10.1126/science.274.5294.1926. url: https://www.science.org/doi/10.1126/science.274.5294.1926.

[9] Arthur S. Reber. “Implicit learning of artificial grammars”. In: Journal of Verbal Learning and Verbal Behavior 6.6 (1967), pp. 855–863. issn: 00225371. doi: 10.1016/S0022-5371(67)80149-X.

[10] Elisabeth A. Karuza et al. “The neural correlates of statistical learning in a word segmentation task: An fMRI study”. In: Brain and Language 127.1 (2013), pp. 46–54. issn: 0093934X. doi: 10.1016/j.bandl.2012.11.007. url: http://dx.doi.org/10.1016/j.bandl.2012.11.007.

[11] Mary Jo Nissen and Peter Bullemer. “Attentional requirements of learning: Evidence from performance measures”. In: Cognitive Psychology 19.1 (Jan. 1987), pp. 1–32. issn: 00100285. doi: 10.1016/0010-0285(87)90002-8. url: https://linkinghub.elsevier.com/retrieve/pii/0010028587900028.

[12] Marvin M. Chun and Yuhong Jiang. “Contextual Cueing: Implicit Learning and Memory of Visual Context Guides Spatial Attention”. In: Cognitive Psychology 36.1 (June 1998), pp. 28–71. issn: 00100285. doi: 10.1006/cogp.1998.0681. url: https://linkinghub.elsevier.com/retrieve/pii/S0010028598906818.

[13] Nicholas B. Turk-Browne et al. “Multidimensional visual statistical learning.” In: Journal of Experimental Psychology: Learning, Memory, and Cognition 34.2 (2008), pp. 399–407. issn: 1939-1285. doi: 10.1037/0278-7393.34.2.399. url: http://doi.apa.org/getdoi.cfm?doi=10.1037/0278-7393.34.2.399.

[14] Hillary Schwarb and Eric H. Schumacher. “Generalized lessons about sequence learning from the study of the serial reaction time task”. In: Advances in Cognitive Psychology 8.2 (2012), pp. 165–178. issn: 18951171. doi: 10.2478/v10053-008-0113-1.

[15] Jonathan Reed and Peder Johnson. “Assessing implicit learning with indirect tests: Determining what is learned about sequence structure.” In: Journal of Experimental Psychology: Learning, Memory, and Cognition 20.3 (1994), pp. 585–594. issn: 1939-1285. doi: 10.1037/0278-7393.20.3.585. url: http://doi.apa.org/getdoi.cfm?doi=10.1037/0278-7393.20.3.585.

[16] Robyn Kim et al. “Testing assumptions of statistical learning: Is it long-term and implicit?” In: Neuroscience Letters 461.2 (2009), pp. 145–149. issn: 03043940. doi: 10.1016/j.neulet.2009.06.030.

[17] Pawel Lewicki, Thomas Hill, and Elizabeth Bizot. “Acquisition of procedural knowledge about a pattern of stimuli that cannot be articulated”. In: Cognitive Psychology 20.1 (Jan. 1988), pp. 24–37. issn: 00100285. doi: 10.1016/0010-0285(88)90023-0. url: https://linkinghub.elsevier.com/retrieve/pii/0010028588900230.

[18] Limor Raviv and Inbal Arnon. “The developmental trajectory of children’s auditory and visual statistical learning abilities: modality-based differences in the effect of age”. In: Developmental Science 21.4 (July 2018), e12593. issn: 1363755X. doi: 10.1111/desc.12593. url: https://onlinelibrary.wiley.com/doi/10.1111/desc.12593.

[19] Jessica F. Schwab et al. “Aging and the statistical learning of grammatical form classes”. In: Psychology and Aging 31.5 (2016), pp. 481–487. issn: 19391498. doi: 10.1037/pag0000110.

[20] Natasha Z. Kirkham, Jonathan A. Slemmer, and Scott P. Johnson. “Visual statistical learning in infancy: evidence for a domain general learning mechanism”. In: Cognition 83.2 (Mar. 2002), B35– B42. issn: 00100277. doi: 10.1016/S0010-0277(02)00004-5. url: https://linkinghub.elsevier.com/retrieve/pii/S0010027702000045.

[21] Karen L. Campbell et al. “Age differences in visual statistical learning.” In: Psychology and Aging 27.3 (Sept. 2012), pp. 650–656. issn: 1939-1498. doi: 10.1037/a0026780. url: http://doi.apa.org/getdoi.cfm?doi=10.1037/a0026780.

[22] Chiara Santolin and Jenny R. Saffran. “Constraints on Statistical Learning Across Species”. In: Trends in Cognitive Sciences 22.1 (Jan. 2018), pp. 52–63. issn: 13646613. doi: 10.1016/j.tics.2017.10.003. url: http://dx.doi.org/10.1016/j.tics.2017.10.003%20https://linkinghub.elsevier.com/retrieve/pii/S1364661317302231.

[23] Juan M. Toro and Josep B. Trobalón. “Statistical computations over a speech stream in a rodent”. In: Perception and Psychophysics 67.5 (2005), pp. 867–875. issn: 00315117. doi: 10.3758/BF03193539.

[24] Toni Cunillera et al. “Time course and functional neuroanatomy of speech segmentation in adults”. In: NeuroImage 48.3 (Nov. 2009), pp. 541–553. issn: 10538119. doi: 10.1016/j.neuroimage.2009.06.069. url: https://linkinghub.elsevier.com/retrieve/pii/S1053811909007095.

[25] Kristin McNealy, John C. Mazziotta, and Mirella Dapretto. “Cracking the language code: Neural mechanisms underlying speech parsing”. In: Journal of Neuroscience 26.29 (2006), pp. 7629–7639. issn: 02706474. doi: 10.1523/JNEUROSCI.5501-05.2006.

[26] Nicholas B. Turk-Browne et al. “Neural Evidence of Statistical Learning: Efficient Detection of Visual Regularities Without Awareness”. In: Journal of Cognitive Neuroscience 21.10 (Oct. 2009), pp. 1934– 1945. issn: 0898-929X. doi: 10.1162/jocn.2009.21131. url: https://direct.mit.edu/jocn/article/21/10/1934/4727/Neural-Evidence-of-Statistical-Learning-Efficient.

[27] Dilshat Abla and Kazuo Okanoya. “Statistical segmentation of tone sequences activates the left inferior frontal cortex: A near-infrared spectroscopy study”. In: Neuropsychologia 46.11 (Sept. 2008), pp. 2787–2795. issn: 00283932. doi: 10.1016/j.neuropsychologia.2008.05.012. url: https://linkinghub.elsevier.com/retrieve/pii/S0028393208002145.

[28] Yasushi Miyashita. “Neuronal correlate of visual associative long-term memory in the primate temporal cortex”. In: Nature (1988), p. 20.

[29] Adam Messinger et al. “Neuronal representations of stimulus associations develop in the temporal lobe during learning”. In: Proceedings of the National Academy of Sciences of the United States of America 98.21 (2001), pp. 12239–12244. issn: 00278424. doi: 10.1073/pnas.211431098.

[30] Cynthia A. Erickson and Robert Desimone. “Responses of macaque perirhinal neurons during and after visual stimulus association learning”. In: Journal of Neuroscience 19.23 (1999), pp. 10404–10416. issn: 02706474. doi: 10.1523/jneurosci.19-23-10404.1999.

[31] Anna C. Schapiro, Lauren V. Kustner, and Nicholas B. Turk-Browne. “Shaping of object representations in the human medial temporal lobe based on temporal regularities”. In: Current Biology 22.17 (2012), pp. 1622–1627. issn: 09609822. doi: 10.1016/j.cub.2012.06.056. url: http://dx.doi.org/10.1016/j.cub.2012.06.056.

[32] Anna C. Schapiro et al. “The Necessity of the Medial Temporal Lobe for Statistical Learning”. In: Journal of Cognitive Neuroscience 26.8 (Aug. 2014), pp. 1736–1747. issn: 0898-929X. doi: 10.1162/jocn_a_00578. url: https://direct.mit.edu/jocn/article/26/8/1736/28163/The-Necessity-of-the-Medial-Temporal-Lobe-for.

[33] Neal J. Cohen and Howard Eichenbaum. Memory, amnesia, and the hippocampal system. Cambridge, Massachusetts: The MIT Press, 1993.

[34] Haline E. Schendan et al. “An fMRI study of the role of the medial temporal lobe in implicit and explicit sequence learning”. In: Neuron 37.6 (2003), pp. 1013–1025. issn: 08966273. doi: 10.1016/S0896-6273(03)00123-5.

[35] Anna C. Schapiro et al. “Neural representations of events arise from temporal community structure”. In: Nature Neuroscience 16.4 (Apr. 2013), pp. 486–492. issn: 1097-6256. doi: 10.1038/nn.3331. url: http://www.nature.com/articles/nn.3331.

[36] Anne G E Collins. “The Cost of Structure Learning”. In: J Cogn Neurosci 29.10 (2017), pp. 1646– 1655.

[37] Ida Momennejad. “Learning Structures: Predictive Representations, Replay, and Generalization”. In: Curr Opin Behav Sci 32 (2020), pp. 155–166.

[38] Christopher W. Lynn and Danielle S. Bassett. “How humans learn and represent networks”. In: Proceedings of the National Academy of Sciences 117.47 (Nov. 2020), pp. 29407–29415. issn: 0027-8424. doi: 10.1073/pnas.1912328117. url: http://www.pnas.org/lookup/doi/10.1073/pnas.1912328117.

[39] Christopher W. Lynn et al. “Human information processing in complex networks”. In: Nature Physics 16.9 (Sept. 2020), pp. 965–973. issn: 1745-2473. doi: 10.1038/s41567-020-0924-7. url: http://www.nature.com/articles/s41567-020-0924-7.

[40] Elisabeth A. Karuza, Sharon L. Thompson-Schill, and Danielle S. Bassett. “Local Patterns to Global Architectures: Influences of Network Topology on Human Learning”. In: Trends in Cognitive Sciences 20.8 (2016), pp. 629–640. issn: 1879307X. doi: 10.1016/j.tics.2016.06.003. url: http://dx.doi.org/10.1016/j.tics.2016.06.003.

[41] Elisabeth A. Karuza et al. “Process reveals structure: How a network is traversed mediates expectations about its architecture”. In: Scientific Reports 7.1 (Dec. 2017), pp. 1–9. issn: 20452322. doi: 10.1038/s41598-017-12876-5. url: http://www.nature.com/articles/s41598-017-12876-5.

[42] Elisabeth A. Karuza, Ari E. Kahn, and Danielle S. Bassett. “Human sensitivity to community structure is robust to topological variation”. In: Complexity 2019 (2019). issn: 10990526. doi: 10.1155/2019/8379321.

[43] Ari E. Kahn et al. “Network constraints on learnability of probabilistic motor sequences”. In: Nature Human Behaviour 2.12 (Dec. 2018), pp. 936–947. issn: 2397-3374. doi: 10.1038/s41562-018-0463-8. url: http://www.nature.com/articles/s41562-018-0463-8.

[44] Steven H. Tompson et al. “Individual differences in learning social and nonsocial network structures.” In: Journal of Experimental Psychology: Learning, Memory, and Cognition 45.2 (Feb. 2019), pp. 253– 271. issn: 1939-1285. doi: 10.1037/xlm0000580. url: http://doi.apa.org/getdoi.cfm?doi=10.1037/xlm0000580.

[45] Anna C. Schapiro et al. “Statistical learning of temporal community structure in the hippocampus”. In: Hippocampus 26.1 (Jan. 2016), pp. 3–8. issn: 10509631. doi: 10.1002/hipo.22523. url: https://onlinelibrary.wiley.com/doi/10.1002/hipo.22523.

[46] Ari E. Kahn et al. “Network structure influences the strength of learned neural representations”. In: Nature Communications 16.1 (Jan. 2025), p. 994. issn: 2041-1723. doi: 10.1038/s41467-024-55459-5. url: https://doi.org/10.1038/s41467-024-55459-5.

[47] Steven H. Tompson et al. “Functional brain network architecture supporting the learning of social networks in humans”. In: NeuroImage 210.January (2020), p. 116498. issn: 10959572. doi: 10.1016/j.neuroimage.2019.116498. url: https://doi.org/10.1016/j.neuroimage.2019.116498.

[48] Jennifer Stiso et al. “Neurophysiological Evidence for Cognitive Map Formation during Sequence Learning”. In: eneuro 9.2 (Mar. 2022), pp. 0361–21. issn: 2373-2822. doi: 10.1523/ENEURO.0361-21.2022. url: https://www.eneuro.org/lookup/doi/10.1523/ENEURO.0361-21.2022.

[49] Laura J. Batterink, Ken A. Paller, and Paul J. Reber. “Understanding the Neural Bases of Implicit and Statistical Learning”. In: Topics in Cognitive Science 11.3 (Apr. 2019), tops.12420. issn: 1756-8757. doi: 10.1111/tops.12420. url: https://onlinelibrary.wiley.com/doi/10.1111/tops.12420.

[50] Danielle S. Bassett et al. “Dynamic reconfiguration of human brain networks during learning”. In: Proceedings of the National Academy of Sciences 108.18 (May 2011), pp. 7641–7646. issn: 0027-8424. doi: 10.1073/pnas.1018985108. url: http://www.pnas.org/cgi/doi/10.1073/pnas.1018985108.

[51] Danielle S. Bassett et al. “Learning-induced autonomy of sensorimotor systems”. In: Nature Neuroscience 18.5 (May 2015), pp. 744–751. issn: 1097-6256. doi: 10.1038/nn.3993. url: http://www.nature.com/articles/nn.3993.

[52] Urs Braun et al. “Dynamic reconfiguration of frontal brain networks during executive cognition in humans”. In: Proceedings of the National Academy of Sciences 112.37 (Sept. 2015), pp. 11678–11683. issn: 0027-8424. doi: 10.1073/pnas.1422487112. url: http://www.pnas.org/lookup/doi/10.1073/pnas.1422487112.

[53] Karolina Finc et al. “Dynamic reconfiguration of functional brain networks during working memory training”. In: Nature Communications 11.1 (Dec. 2020), p. 2435. doi: 10.1038/s41467-020-15631-z. url: http://www.nature.com/articles/s41467-020-15631-z.

[54] Alexander Schaefer et al. “Local-Global Parcellation of the Human Cerebral Cortex from Intrinsic Functional Connectivity MRI”. In: Cerebral Cortex 28.9 (Sept. 2018), pp. 3095–3114. issn: 1047-3211. doi: 10.1093/cercor/bhx179. url: https://academic.oup.com/cercor/article/28/9/3095/3978804.

[55] Randy L. Buckner et al. “The organization of the human cerebellum estimated by intrinsic functional connectivity”. In: Journal of Neurophysiology 106.5 (Nov. 2011), pp. 2322–2345. issn: 0022-3077. doi: 10.1152/jn.00339.2011. url: https://www.physiology.org/doi/10.1152/jn.00339.2011.

[56] Ye Tian et al. “Topographic organization of the human subcortex unveiled with functional connectivity gradients”. In: Nature Neuroscience 23.11 (2020), pp. 1421–1432. issn: 15461726. doi: 10.1038/s41593-020-00711-6. url: http://dx.doi.org/10.1038/s41593-020-00711-6.

[57] Danielle S. Bassett et al. “Task-Based Core-Periphery Organization of Human Brain Dynamics”. In: PLoS Computational Biology 9.9 (Sept. 2013). Ed. by Olaf Sporns, e1003171. issn: 1553-7358. doi: 10.1371/journal.pcbi.1003171. url: https://dx.plos.org/10.1371/journal.pcbi.1003171.

[58] Marcelo G. Mattar et al. “Predicting future learning from baseline network architecture”. In: NeuroImage 172.January (2018), pp. 107–117. issn: 10959572. doi: 10.1016/j.neuroimage.2018.01.037. url: https://doi.org/10.1016/j.neuroimage.2018.01.037.

[59] Javier Gonzalez-Castillo and Peter A. Bandettini. “Task-based dynamic functional connectivity: Recent findings and open questions”. In: NeuroImage 180.August 2017 (2018), pp. 526–533. issn: 10959572. doi: 10.1016/j.neuroimage.2017.08.006. url: https://doi.org/10.1016/j.neuroimage.2017.08.006.

[60] James M. Shine et al. “Estimation of dynamic functional connectivity using Multiplication of Temporal Derivatives”. In: NeuroImage 122 (2015), pp. 399–407. issn: 10959572. doi: 10.1016/j.neuroimage.2015.07.064. url: http://dx.doi.org/10.1016/j.neuroimage.2015.07.064.

[61] Lucy R. Chai et al. “Functional Network Dynamics of the Language System”. In: Cerebral Cortex 26.11 (Oct. 2016), pp. 4148–4159. issn: 1047-3211. doi: 10.1093/cercor/bhw238. url: https://academic.oup.com/cercor/article-lookup/doi/10.1093/cercor/bhw238.

[62] Soyoung Kwon et al. “Attention reorganizes connectivity across networks in a frequency specific manner”. In: NeuroImage 144.March 2016 (2017), pp. 217–226. issn: 10959572. doi: 10.1016/j.neuroimage.2016.10.014. url: http://dx.doi.org/10.1016/j.neuroimage.2016.10.014.

[63] Sara Spadone et al. “Dynamic reorganization of human resting-state networks during visuospatial attention”. In: Proceedings of the National Academy of Sciences of the United States of America 112.26 (2015), pp. 8112–8117. issn: 10916490. doi: 10.1073/pnas.1415439112.

[64] Marcelo G. Mattar et al. “A Functional Cartography of Cognitive Systems”. In: PLOS Computational Biology 11.12 (Dec. 2015). Ed. by Christopher J Honey, e1004533. issn: 1553-7358. doi: 10.1371/journal.pcbi.1004533. url: https://dx.plos.org/10.1371/journal.pcbi.1004533.

[65] James M. Shine and Russell A. Poldrack. “Principles of dynamic network reconfiguration across diverse brain states”. In: NeuroImage 180.July 2017 (2018), pp. 396–405. issn: 10959572. doi: 10.1016/j.neuroimage.2017.08.010. url: https://doi.org/10.1016/j.neuroimage.2017.08.010.

[66] R. Nathan Spreng et al. “Default network activity, coupled with the frontoparietal control network, supports goal-directed cognition”. In: NeuroImage 53.1 (2010), pp. 303–317. issn: 10538119. doi: 10.1016/j.neuroimage.2010.06.016. url: http://dx.doi.org/10.1016/j.neuroimage.2010.06.016.

[67] Bastien Herlin, Vincent Navarro, and Sophie Dupont. “The temporal pole: From anatomy to function—A literature appraisal”. In: Journal of Chemical Neuroanatomy 113.January (Apr. 2021), p. 101925. issn: 08910618. doi: 10.1016/j.jchemneu.2021.101925. url: https://linkinghub.elsevier.com/retrieve/pii/S0891061821000089.

[68] Roel M. Willems et al. “Prediction during Natural Language Comprehension”. In: Cerebral Cortex 26.6 (June 2016), pp. 2506–2516. issn: 14602199. doi: 10.1093/cercor/bhv075. url: https://academic.oup.com/cercor/article-lookup/doi/10.1093/cercor/bhv075.

[69] Samuel Nastase, Vittorio Iacovella, and Uri Hasson. “Uncertainty in visual and auditory series is coded by modality-general and modality-specific neural systems”. In: Human Brain Mapping 35.4 (2014), pp. 1111–1128. issn: 10970193. doi: 10.1002/hbm.22238.

[70] Iris Vilares et al. “Differential Representations of Prior and Likelihood Uncertainty in the Human Brain”. In: Current Biology 22.18 (Sept. 2012), pp. 1641–1648. doi: 10.1016/j.cub.2012.07.010. url: https://linkinghub.elsevier.com/retrieve/pii/S0960982212008019.

[71] Nicholas L. Balderston, Doug H. Schultz, and Fred J. Helmstetter. “The human amygdala plays a stimulus specific role in the detection of novelty”. In: NeuroImage 55.4 (2011), pp. 1889–1898. issn: 10538119. doi: 10.1016/j.neuroimage.2011.01.034. url: http://dx.doi.org/10.1016/j.neuroimage.2011.01.034.

[72] Edmund T. Rolls et al. “The human posterior cingulate, retrosplenial, and medial parietal cortex effective connectome, and implications for memory and navigation”. In: Human Brain Mapping 44.2 (2023), pp. 629–655. issn: 10970193. doi: 10.1002/hbm.26089.

[73] Edmund T. Rolls et al. “Multiple cortical visual streams in humans”. In: Cerebral Cortex 33.7 (2023), pp. 3319–3349. issn: 14602199. doi: 10.1093/cercor/bhac276.

[74] Alexandra O. Constantinescu, Jill X. O’Reilly, and Timothy E.J. Behrens. “Organizing conceptual knowledge in humans with a gridlike code”. In: Science 352.6292 (June 2016), pp. 1464–1468. issn: 10959203. doi: 10.1126/science.aaf0941. url: https://www.science.org/doi/10.1126/science.aaf0941.

[75] John O’Keefe and Lynn Nadel. The Hippocampus as a Cognitive Map. Oxford University Press, 1978. isbn: 0198572069. doi: 10.1176/ajp.136.10.1353.

[76] Timothy E.J. Behrens et al. “What Is a Cognitive Map? Organizing Knowledge for Flexible Behavior”. In: Neuron 100.2 (2018), pp. 490–509. issn: 10974199. doi: 10.1016/j.neuron.2018.10.002. url: https://doi.org/10.1016/j.neuron.2018.10.002.

[77] Anna C. Schapiro et al. “Complementary learning systems within the hippocampus: a neural network modelling approach to reconciling episodic memory with statistical learning”. In: Philosophical Transactions of the Royal Society B: Biological Sciences 372.1711 (Jan. 2017), p. 20160049. issn: 0962-8436. doi: 10.1098/rstb.2016.0049. url: https://royalsocietypublishing.org/doi/10.1098/rstb.2016.0049.

[78] L.M. Harrison, A. Duggins, and K.J. Friston. “Encoding uncertainty in the hippocampus”. In: Neural Networks 19.5 (June 2006), pp. 535–546. issn: 08936080. doi: 10.1016/j.neunet.2005.11.002. url: https://linkinghub.elsevier.com/retrieve/pii/S0893608006000025.

[79] William Qian et al. “Optimizing the human learnability of abstract network representations”. In: Proceedings of the National Academy of Sciences of the United States of America 119.35 (2022), pp. 1–10. issn: 10916490. doi: 10.1073/pnas.2121338119.

[80] Catherine W. Tallman, Robert E. Clark, and Christine N. Smith. “Human brain activity and functional connectivity as memories age from one hour to one month”. In: Cognitive Neuroscience 13.3-4 (2022), pp. 115–133. issn: 17588936. doi: 10.1080/17588928.2021.2021164. url: https://doi.org/10.1080/17588928.2021.2021164.

[81] Emily T. Cowan et al. “Memory consolidation as an adaptive process”. In: Psychonomic Bulletin and Review 28.6 (2021), pp. 1796–1810. issn: 15315320. doi: 10.3758/s13423-021-01978-x.

[82] Alexa Tompary and Lila Davachi. “Consolidation Promotes the Emergence of Representational Overlap in the Hippocampus and Medial Prefrontal Cortex”. In: Neuron 96.1 (2017), pp. 228–241. issn: 10974199. doi: 10.1016/j.neuron.2017.09.005. url: https://doi.org/10.1016/j.neuron.2017.09.005.

[83] Simon J. Durrant et al. “Sleep-dependent consolidation of statistical learning”. In: Neuropsychologia 49.5 (2011), pp. 1322–1331. issn: 00283932. doi: 10.1016/j.neuropsychologia.2011.02.015. url: http://dx.doi.org/10.1016/j.neuropsychologia.2011.02.015.

[84] K.J Friston et al. “Psychophysiological and Modulatory Interactions in Neuroimaging”. In: NeuroImage 6.3 (Oct. 1997), pp. 218–229. issn: 10538119. doi: 10.1006/nimg.1997.0291. url: https://linkinghub.elsevier.com/retrieve/pii/S1053811997902913.

[85] B.J. Casey et al. “The Adolescent Brain Cognitive Development (ABCD) study: Imaging acquisition across 21 sites”. In: Developmental Cognitive Neuroscience 32.March (Aug. 2018), pp. 43–54. issn: 18789293. doi: 10.1016/j.dcn.2018.03.001. url: https://doi.org/10.1016/j.dcn.2018.03.001%20https://linkinghub.elsevier.com/retrieve/pii/S1878929317301214.

[86] André J.W. van der Kouwe et al. “Brain morphometry with multiecho MPRAGE”. In: NeuroImage 40.2 (Apr. 2008), pp. 559–569. issn: 10538119. doi: 10.1016/j.neuroimage.2007.12.025. url: https://linkinghub.elsevier.com/retrieve/pii/S1053811907011457.

[87] Mark A. Griswold et al. “Generalized autocalibrating partially parallel acquisitions (GRAPPA)”. In: Magnetic Resonance in Medicine 47.6 (June 2002), pp. 1202–1210. issn: 0740-3194. doi: 10.1002/mrm.10171. url: https://onlinelibrary.wiley.com/doi/10.1002/mrm.10171.

[88] Oscar Esteban et al. “fMRIPrep: a robust preprocessing pipeline for functional MRI”. In: Nature Methods (2018). doi: 10.1038/s41592-018-0235-4.

[89] Oscar Esteban et al. “fMRIPrep”. In: Software (2018). doi: 10.5281/zenodo.852659.

[90] K. Gorgolewski et al. “Nipype: a flexible, lightweight and extensible neuroimaging data processing framework in Python”. In: Frontiers in Neuroinformatics 5 (2011), p. 13. doi: 10.3389/fninf.2011.00013.

[91] Krzysztof J. Gorgolewski et al. “Nipype”. In: Software (2018). doi: 10.5281/zenodo.596855.

[92] N. J. Tustison et al. “N4ITK: Improved N3 Bias Correction”. In: IEEE Transactions on Medical Imaging 29.6 (2010), pp. 1310–1320. issn: 0278-0062. doi: 10.1109/TMI.2010.2046908.

[93] B.B. Avants et al. “Symmetric diffeomorphic image registration with cross-correlation: Evaluating automated labeling of elderly and neurodegenerative brain”. In: Medical Image Analysis 12.1 (2008), pp. 26–41. issn: 1361-8415. doi: 10.1016/j.media.2007.06.004. url: http://www.sciencedirect.com/science/article/pii/S1361841507000606.

[94] Y. Zhang, M. Brady, and S. Smith. “Segmentation of brain MR images through a hidden Markov random field model and the expectation-maximization algorithm”. In: IEEE Transactions on Medical Imaging 20.1 (2001), pp. 45–57. issn: 0278-0062. doi: 10.1109/42.906424.

[95] Anders M. Dale, Bruce Fischl, and Martin I. Sereno. “Cortical Surface-Based Analysis: I. Segmentation and Surface Reconstruction”. In: NeuroImage 9.2 (1999), pp. 179–194. issn: 1053-8119. doi: 10.1006/nimg.1998.0395. url: http://www.sciencedirect.com/science/article/pii/S1053811998903950.

[96] Arno Klein et al. “Mindboggling morphometry of human brains”. In: PLOS Computational Biology 13.2 (2017), e1005350. issn: 1553-7358. doi: 10.1371/journal.pcbi.1005350. url: http://journals.plos.org/ploscompbiol/article?id=10.1371/journal.pcbi.1005350.

[97] VS Fonov et al. “Unbiased nonlinear average age-appropriate brain templates from birth to adulthood”. In: NeuroImage 47, Supplement 1 (2009), S102. doi: 10.1016/S1053-8119(09)70884-5.

[98] AC Evans et al. “Brain templates and atlases”. In: NeuroImage 62.2 (2012), pp. 911–922. doi: 10.1016/j.neuroimage.2012.01.024.

[99] Mark Jenkinson et al. “Improved Optimization for the Robust and Accurate Linear Registration and Motion Correction of Brain Images”. In: NeuroImage 17.2 (2002), pp. 825–841. issn: 1053-8119. doi: 10.1006/nimg.2002.1132. url: http://www.sciencedirect.com/science/article/pii/S1053811902911328.

[100] Robert W. Cox and James S. Hyde. “Software tools for analysis and visualization of fMRI data”. In: NMR in Biomedicine 10.4-5 (1997), pp. 171–178. doi: 10.1002/(SICI)1099-1492(199706/08)10:4/5<171::AID-NBM453>3.0.CO;2-L.

[101] Douglas N Greve and Bruce Fischl. “Accurate and robust brain image alignment using boundary-based registration”. In: NeuroImage 48.1 (2009), pp. 63–72. issn: 1095-9572. doi: 10.1016/j.neuroimage.2009.06.060.

[102] Jonathan D. Power et al. “Methods to detect, characterize, and remove motion artifact in resting state fMRI”. In: NeuroImage 84.Supplement C (2014), pp. 320–341. issn: 1053-8119. doi: 10.1016/j.neuroimage.2013.08.048. url: http://www.sciencedirect.com/science/article/pii/S1053811913009117.

[103] Yashar Behzadi et al. “A component based noise correction method (CompCor) for BOLD and perfusion based fMRI”. In: NeuroImage 37.1 (2007), pp. 90–101. issn: 1053-8119. doi: 10.1016/j.neuroimage.2007.04.042. url: http://www.sciencedirect.com/science/article/pii/S1053811907003837.

[104] Theodore D. Satterthwaite et al. “An improved framework for confound regression and filtering for control of motion artifact in the preprocessing of resting-state functional connectivity data”. In: NeuroImage 64.1 (2013), pp. 240–256. issn: 10538119. doi: 10.1016/j.neuroimage.2012.08.052. url: http://linkinghub.elsevier.com/retrieve/pii/S1053811912008609.

[105] C. Lanczos. “Evaluation of Noisy Data”. In: Journal of the Society for Industrial and Applied Mathematics Series B Numerical Analysis 1.1 (1964), pp. 76–85. issn: 0887-459X. doi: 10.1137/0701007. url: http://epubs.siam.org/doi/10.1137/0701007.

[106] Alexandre Abraham et al. “Machine learning for neuroimaging with scikit-learn”. English. In: Frontiers in Neuroinformatics 8 (2014). issn: 1662-5196. doi: 10.3389/fninf.2014.00014. url: https://www.frontiersin.org/articles/10.3389/fninf.2014.00014/full.

[107] Andrew C. Murphy et al. “Multimodal network dynamics underpinning working memory”. In: Nature Communications 11.1 (Dec. 2020), p. 3035. issn: 2041-1723. doi: 10.1038/s41467-020-15541-0. url: http://www.nature.com/articles/s41467-020-15541-0.

[108] C. M. A. Pennartz et al. “Corticostriatal Interactions during Learning, Memory Processing, and Decision Making”. In: Journal of Neuroscience 29.41 (Oct. 2009), pp. 12831–12838. issn: 0270-6474. doi: 10.1523/JNEUROSCI.3177-09.2009. url: https://www.jneurosci.org/lookup/doi/10.1523/JNEUROSCI.3177-09.2009.

[109] Okihide Hikosaka et al. “Central mechanisms of motor skill learning”. In: Current Opinion in Neurobiology 12.2 (Apr. 2002), pp. 217–222. issn: 09594388. doi: 10.1016/S0959-4388(02)00307-0. url: https://linkinghub.elsevier.com/retrieve/pii/S0959438802003070.

[110] Eran Dayan and Leonardo G. Cohen. “Neuroplasticity Subserving Motor Skill Learning”. In: Neuron 72.3 (Nov. 2011), pp. 443–454. issn: 08966273. doi: 10.1016/j.neuron.2011.10.008. url: http://dx.doi.org/10.1016/j.neuron.2011.10.008%20 https://linkinghub.elsevier.com/retrieve/pii/S0896627311009184.

[111] Karl J. Friston. “Functional and Effective Connectivity: A Review”. In: Brain Connectivity 1.1 (Jan. 2011), pp. 13–36. issn: 2158-0014. doi: 10.1089/brain.2011.0008. url: http://www.liebertpub.com/doi/10.1089/brain.2011.0008.

[112] Arun S. Mahadevan et al. “Evaluating the sensitivity of functional connectivity measures to motion artifact in resting-state fMRI data”. In: NeuroImage 241.July (Nov. 2021), p. 118408. issn: 10538119. doi: 10.1016/j.neuroimage.2021.118408. url: https://doi.org/10.1016/j.neuroimage.2021.118408%20https://linkinghub.elsevier.com/retrieve/pii/S1053811921006832.

[113] Andrew Zalesky, Alex Fornito, and Ed Bullmore. “On the use of correlation as a measure of network connectivity”. In: NeuroImage 60.4 (May 2012), pp. 2096–2106. issn: 10538119. doi: 10.1016/j.neuroimage.2012.02.001. url: http://dx.doi.org/10.1016/j.neuroimage.2012.02.001%20https://linkinghub.elsevier.com/retrieve/pii/S1053811912001784.

[114] Catie Chang and Gary H. Glover. “Time-frequency dynamics of resting-state brain connectivity measured with fMRI”. In: NeuroImage 50.1 (2010), pp. 81–98. issn: 10538119. doi: 10.1016/j.neuroimage.2009.12.011. url: http://dx.doi.org/10.1016/j.neuroimage.2009.12.011.

[115] Ünal Sakoğlu et al. “A method for evaluating dynamic functional network connectivity and taskmodulation: application to schizophrenia”. In: Magnetic Resonance Materials in Physics, Biology and Medicine 23.5-6 (Dec. 2010), pp. 351–366. issn: 0968-5243. doi: 10.1007/s10334-010-0197-8. url: http://link.springer.com/10.1007/s10334-010-0197-8.

[116] R. Matthew Hutchison et al. “Dynamic functional connectivity: Promise, issues, and interpretations”. In: NeuroImage 80 (2013), pp. 360–378. issn: 10538119. doi: 10.1016/j.neuroimage.2013.05.079. url: http://dx.doi.org/10.1016/j.neuroimage.2013.05.079.

[117] Nora Leonardi and Dimitri Van De Ville. “On spurious and real fluctuations of dynamic functional connectivity during rest”. In: NeuroImage 104 (2015), pp. 430–436. issn: 10959572. doi: 10.1016/j.neuroimage.2014.09.007. url: http://dx.doi.org/10.1016/j.neuroimage.2014.09.007.

[118] R. Matthew Hutchison et al. “Resting-state networks show dynamic functional connectivity in awake humans and anesthetized macaques”. In: Human Brain Mapping 34.9 (2013), pp. 2154–2177. issn: 10659471. doi: 10.1002/hbm.22058.

[119] Maxwell A. Bertolero et al. “A mechanistic model of connector hubs, modularity and cognition”. In: Nature Human Behaviour 2.10 (Oct. 2018), pp. 765–777. issn: 2397-3374. doi: 10.1038/s41562-018-0420-6. url: http://www.nature.com/articles/s41562-018-0420-6.

[120] Mark Newman. Networks. Oxford University Press, Mar. 2010. isbn: 9780199206650. doi: 10.1093/acprof:oso/9780199206650.001.0001. url: https://oxford.universitypressscholarship.com/view/10.1093/acprof:oso/9780199206650.001.0001/acprof-9780199206650.

[121] Peter J. Mucha et al. “Community Structure in Time-Dependent, Multiscale, and Multiplex Networks”. In: Science 328.5980 (May 2010), pp. 876–878. issn: 0036-8075. doi: 10.1126/science.1184819. url: https://www.science.org/doi/10.1126/science.1184819%20https://www.sciencemag.org/lookup/doi/10.1126/science.1184819.

[122] Benjamin H. Good, Yves-Alexandre de Montjoye, and Aaron Clauset. “Performance of modularity maximization in practical contexts”. In: Physical Review E 81.4 (Apr. 2010), p. 046106. issn: 15393755. doi: 10.1103/PhysRevE.81.046106. url: https://link.aps.org/doi/10.1103/PhysRevE.81.046106.

[123] David M. Lydon-Staley et al. “Evaluation of confound regression strategies for the mitigation of micromovement artifact in studies of dynamic resting-state functional connectivity and multilayer network modularity”. In: Network Neuroscience 3.2 (Jan. 2019), pp. 427–454. issn: 2472-1751. doi: 10.1162/netn_a_00071. url: https://direct.mit.edu/netn/article/3/2/427-454/2228.

[124] Alexandra Kuznetsova, Per B. Brockhoff, and Rune H. B. Christensen. “lmerTest Package: Tests in Linear Mixed Effects Models”. In: Journal of Statistical Software 82.13 (2017). issn: 1548-7660. doi: 10.18637/jss.v082.i13. url: http://www.jstatsoft.org/v82/i13/.

[125] Yoav Benjamini and Yosef Hochberg. “Controlling the False Discovery Rate: A Practical and Powerful Approach to Multiple Testing”. In: Journal of the Royal Statistical Society: Series B (Methodological) 57.1 (Jan. 1995), pp. 289–300. issn: 00359246. doi: 10.1111/j.2517-6161.1995.tb02031.x. url: https://onlinelibrary.wiley.com/doi/10.1111/j.2517-6161.1995.tb02031.x.

[126] Sara McLaughlin Mitchell, Samantha Lange, and Holly Brus. “Gendered citation patterns in international relations journals”. In: International Studies Perspectives 14.4 (2013), pp. 485–492.

[127] Michelle L Dion, Jane Lawrence Sumner, and Sara McLaughlin Mitchell. “Gendered citation patterns across political science and social science methodology fields”. In: Political Analysis 26.3 (2018), pp. 312–327.

[128] Neven Caplar, Sandro Tacchella, and Simon Birrer. “Quantitative evaluation of gender bias in astronomical publications from citation counts”. In: Nature Astronomy 1.6 (2017), p. 0141.

[129] Daniel Maliniak, Ryan Powers, and Barbara F Walter. “The gender citation gap in international relations”. In: International Organization 67.4 (2013), pp. 889–922.

[130] Jordan D. Dworkin et al. “The extent and drivers of gender imbalance in neuroscience reference lists”. In: bioRxiv (2020). doi: 10.1101/2020.01.03.894378. eprint: https://www.biorxiv.org/content/early/2020/01/11/2020.01.03.894378.full.pdf. xurl: https://www.biorxiv.org/content/early/2020/01/11/2020.01.03.894378.

[131] Maxwell A. Bertolero et al. “Racial and ethnic imbalance in neuroscience reference lists and intersections with gender”. In: bioRxiv (2020).

[132] Xinyi Wang et al. “Gendered citation practices in the field of communication”. In: Annals of the International Communication Association (2021). doi: 10.1080/23808985.2021.1960180.

[133] Paula Chatterjee and Rachel M Werner. “Gender Disparity in Citations in High-Impact Journal Articles”. In: JAMA Netw Open 4.7 (2021), e2114509.

[134] Jacqueline M Fulvio, Ileri Akinnola, and Bradley R Postle. “Gender (Im)balance in Citation Practices in Cognitive Neuroscience”. In: J Cogn Neurosci 33.1 (2021), pp. 3–7.

[135] Dale Zhou et al. Gender Diversity Statement and Code Notebook v1.0. Version v1.0. Feb. 2020. doi: 10.5281/zenodo.3672110. url: https://doi.org/10.5281/zenodo.3672110.

[136] Anurag Ambekar et al. “Name-ethnicity classification from open sources”. In: Proceedings of the 15th ACM SIGKDD international conference on Knowledge Discovery and Data Mining. 2009, pp. 49–58.

[137] Gaurav Sood and Suriyan Laohaprapanon. “Predicting race and ethnicity from the sequence of characters in a name”. In: arXiv preprint 1805.02109 (2018).

